# CAR T cell therapy drives endogenous locoregional T cell dynamics in a responding patient with glioblastoma

**DOI:** 10.1101/2021.09.22.460392

**Authors:** Vanessa D. Jonsson, Rachel H. Ng, Natalie Dullerud, Robyn A. Wong, Jonathan Hibbard, Dongrui Wang, Brenda Aguilar, Renate Starr, Lihong Weng, Darya Alizadeh, Stephen J. Forman, Behnam Badie, Christine E. Brown

## Abstract

CAR T cell therapy has transformed clinical care and management of patients with certain hematological cancers. However, it remains unclear whether the success of CAR T cell therapy relies solely on CAR T cell engagement with tumor antigen, or if it also requires the stimulation of an individual patient’s endogenous T cell response. Here, we performed combined analysis of longitudinal, single cell RNA and T cell receptor sequencing on glioblastoma tumors, peripheral blood (PB), and cerebrospinal fluid (CSF) from a patient with recurrent multifocal glioblastoma that underwent a remarkable response followed by recurrence on IL13RA2-targeted CAR T cell therapy (Brown et al. 2016). Single cell analysis of a tumor resected prior to CAR T cell therapy revealed the existence of an inflamed tumor microenvironment including a CD8+ cytotoxic, clonally expanded and antigen specific T cell population that disappeared in the recurrent setting. Longitudinal tracking of T cell receptors uncovered distinct T cell dynamics classes in the CSF during CAR T cell therapy. These included T cell clones with transient dynamics, representing intraventricular CAR T cell delivery and endogenous T cell recruitment from the PB into the CSF; and a group of T cells in the cerebrospinal fluid, that tracked with clonally expanded tumor resident T cells and whose dynamics contracted concomitantly with tumor volume. Our results suggest the existence of an endogenous T cell population that was invigorated by intraventricular CAR T cell infusions, and combined with the therapy to produce a complete response.

---

We previously reported on a patient with recurrent multifocal glioblastoma, who was treated with CAR T cells targeting tumor- associated antigen interleukin-13 receptor alpha 2 (IL13Rα2) and who sustained a complete response despite non-uniform antigen expression and after 7.5 months recurred with tumors arising at new sites (Brown et al. 2016). This patient’s disease course— a remarkable regression of all brain and spinal tumors following CAR T cell therapy— provided the opportunity to characterize cancer immune dynamics in one patient and two clinical settings — during response and recurrence (Fig. 1a). Here, we performed combined immunogenomics analyses on single cell, bulk RNA and T cell receptor (TCR) sequencing data from this patient’s tumor, CAR T cell infusion products, cerebrospinal fluid (CSF), and peripheral blood (PB) samples collected longitudinally over the course of CAR T cell therapy.

**Fig. 1.**
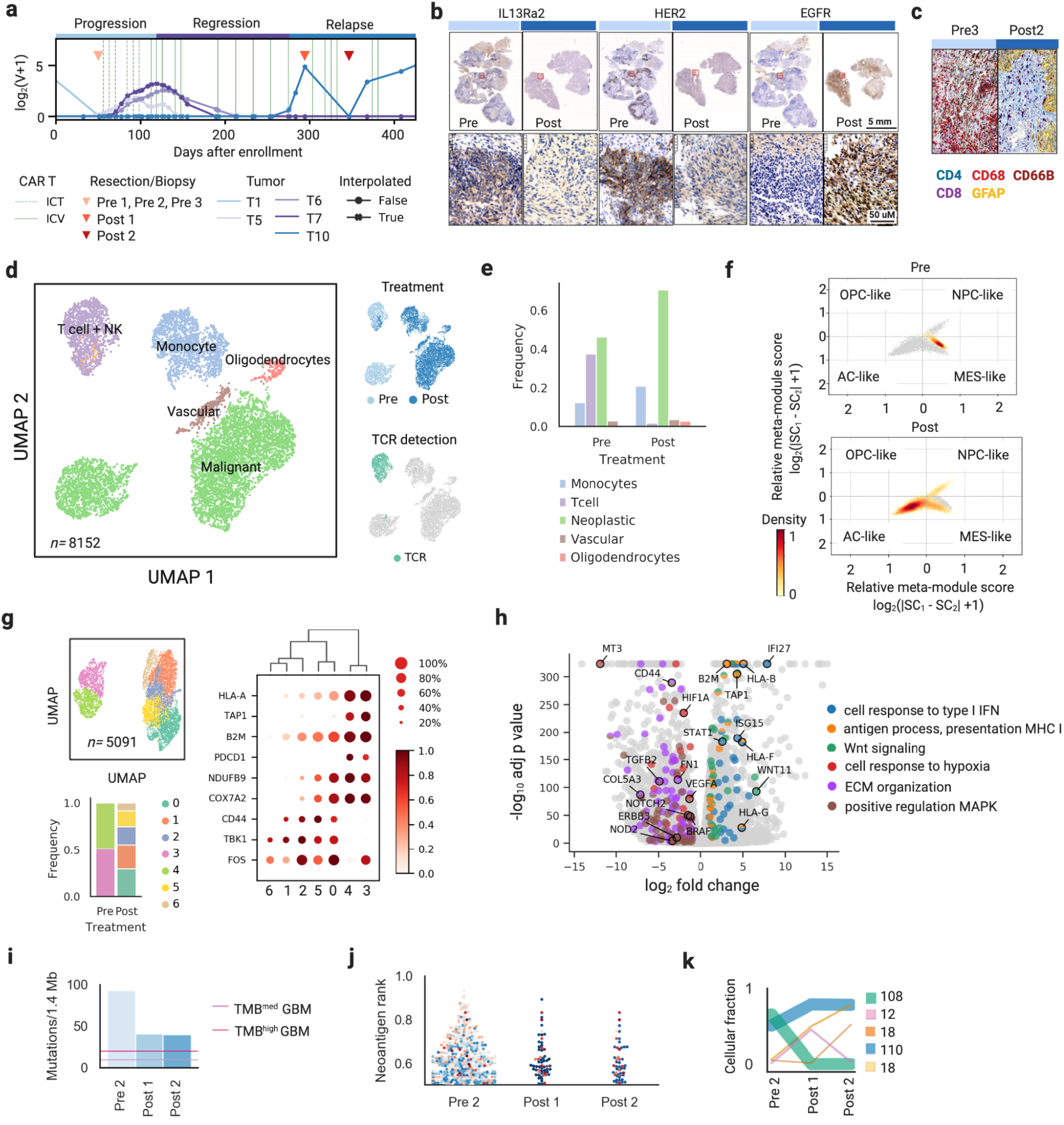
Characterization of pre-therapy and recurrent GBM tumors in a patient that received IL13Ra2 CAR T cells for recurrent glioblastoma. Pre-treatment tumors (Pre1, Pre2, Pre3) and post-treatment tumors (Post1, Post2) are characterized by bulk RNA and exome sequencing. Single cell RNA sequencing performed on Pre1 and Post2 tumors, labeled as Pre and Post respectively. a, Clinical course, measured an interpolated tumor volume over the course of CAR T cell therapy of a glioblastoma patient reported in Brown et al, 2017 (Methods). b, Immunohistochemistry evaluation of GBM surface antigen expression of IL13Ra2, HER2 and EGFR. c, Chromogen staining of CD4 CD8, CD68, GFAP and CD66B surface antigen markers. d, Uniform manifold approximation and projection (UMAP) of all tumor-resident cells before and after treatment. Clusters are labelled by inferred cell types – neoplastic cells, T cells, NK cells, monocytes, oligodendrocytes, and vascular cells (left). UMAP is colored by treatment status (top right), and TCR detection (bottom right). e, Frequency bar plot of cell types per treatment status. f, Two-dimensional representation of GBM cellular state colored by cell density per treatment status; each quadrant corresponds to a relative score for one of four cellular states (Mesenchymal/MES-like, Astrocytic/AS-like, Oligodendrocytic/OPC-like, and Neural progenitor/NPC-like) derived in (Suva). g, UMAP of pre- and post-treatment malignant cell clusters (top left). Frequency of each malignant cell cluster in each tumor (bottom left). Dot plot of genes differentially expressed in malignant clusters, where color represents average expression and size represents percentage of cells expressing. h, Volcano plot of differential expression results comparing pre- and post-treatment malignant cells. Selected GO Biological Processes significantly upregulated in each condition are colored and annotated by representative genes. i, Summary of mutation burden detected in whole exome sequencing. Lines indicate lower bounds for medium and high mutation burden in glioma. j, Swarm plot of neoantigen burden pre-therapy and during recurrence based on exome sequencing; variants were scored with respect to whether the peptide was found to bind to the MHC allele with a binding strength of less than 500 nM and its wild- type cognate peptide bound to the same allele with a binding strength of greater than 500 nM. k, Changes in clonal mutation composition detected in exome sequencing in pre-treated and recurrent tumors; persistent mutations are in blue, mutation clusters with decreasing cellular prevalence are in green, and increasing in cellular prevalence are in orange and pink (left).

## RESULTS

### Changes in tumor antigen expression and GBM cellular phenotype following CAR T cell therapy

Immunohistochemistry of tumor associated antigens (IL13Rα2, HER2, and EGFR) on pre- and post-CAR T cell treated tumors revealed a reduction in IL13Rα2 and HER2, and a gain in EGFR expression, and highlighted the diversity of glioma associated antigen surface expression characteristic in GBM tumors (Schäfer et al. 2019). This supported a response to IL13BBζ-CAR T cells targeting of antigen-positive populations with the ultimate emergence of antigen-low or -negative tumor relapse (Fig. 1b). Simultaneously, chromogen staining of immune cell markers (GFAP, CD4, CD8, and CD68) uncovered changes in the tumor immune landscape, including a loss of CD8+ T cell infiltrates at relapse (Fig. 1c).

We generated 5′ single cell RNA sequencing (scRNA seq) and T cell receptor sequencing (TCR seq) libraries from two tumors for this patient – one pre-treatment tumor (Pre 1) and a second tumor (Post 2), harvested during disease recurrence (Fig. 1c) and retained cell transcriptomes from a total of 8152 cells, with paired TCR sequences in 1015 out of 1205 T cells (84%; Fig. 1d, Extended Data Fig. 2a-d, Methods). We identified six cell clusters including two malignant/neoplastic clusters, and one cluster each for monocytes, T cells (clustered with NK cells), vascular cells, and oligodendrocytes. Differences in the relative frequencies of malignant and immune cell types were discovered in pre and post treatment tumors: a reduction of T cells (37.6% Pre, 1.8% Post) and increases of monocytes (12.3% Pre, 21.1% Post), and malignant cells (46.3% Pre, 70.4% Post) (Fig. 1e).

To explore tumor intrinsic genetic factors that could modulate antigen expression and the tumor immune environment, we focused our analysis on single cell RNA seq gene expression profiles of 5091 malignant cells (Fig. 1g). Meta module analysis of malignant cells identified changes in GBM cellular phenotypes in pre- and post- treatment tumors (Methods); including a transition from a mesenchymal-like (MES) phenotype, to a mixed MES-like, neural- progenitor-like (NPC) and astrocyte-like (AC) phenotype (Fig. 1f, Extended Data Fig. 1c). We observed an increase in EGFR surface antigen expression (log2(FC(Post/Pre))=1.05) associated with the AC-like post-treatment tumor (Extended Data Fig. 2e), consistent with results demonstrating that the AC-like GBM phenotype is driven by high EGFR amplification (Verhaak et al. 2010; Neftel et al. 2019), while IL13Ra2 expression is associated with a mesenchymal gene expression (Brown et al. 2013). Gene fusion analysis on bulk RNA sequencing in pre and post treatment tumors confirmed the presence of FGFR3-TACC3 oncogenic driver (Singh et al. 2012), with amplification of this gene fusion increasing at relapse (Extended Data Fig. 1b).

Combined analysis demonstrated this patient’s malignant cells exhibited mixed GBM cellular phenotypes, heterogeneous tumor antigen expression and gene expression that clustered entirely by treatment status (Fig. 1d,g, Extended Data Fig. 1a): this substantiates existing results that demonstrate intra-patient and intra-tumoral diversity present in GBM. Moreover, changes in GBM phenotype and tumor associated antigen expression suggest that CAR T cells targeted IL13Rα2 on MES-like tumors, which was followed by recurrence of EGFR positive tumor dominant in AC-like cells; and highlight a possible role of CAR T cells in sculpting GBM cellular phenotypes through antigen targeting.

### Tumor transcriptional changes reveal a transition in the TME from inflamed to immune excluded

We then investigated tumor intrinsic factors that could modulate the tumor immune environment. We performed differential gene expression analysis of pre- and post-treatment malignant single cells — genes upregulated in pre-treatment malignant cells were enriched in major histocompatibility complex (MHC) class I antigen processing and presentation (TAP1, B2M, HLA; Adjusted p-value= 4.8 × 10-11) and type I IFN response (IRF1, STAT1; Adjusted p-value = 2.4×10-11), suggesting that although this patient was rapidly progressing prior to the initiation of CAR therapy there was a pre-existing antitumor immune reactivity, with induced interferon-γ (IFNγ) signaling and subsequent PD-L1 upregulation (Adjusted p-value = 1.0 × 10-11) (Fig. 1h, Extended Data Fig. 1d). Furthermore, gene expression pathways associated with reduced immune cell infiltration and immunosuppression, namely extracellular matrix (ECM) organization (COL5A3, FN1; Adjusted p-value = 4.1 × 10-11), hypoxia response (HIF1A Adjusted p-value = 2.1 × 10-2), were upregulated in the recurrent tumor. Somatic variant calling on pre-treatment tumor whole exome sequencing revealed a high tumor mutation burden (TMB) (93 mutations/1.4 Mbp) for GBM (Hodges et al. 2017) (Fig. 1i). Analysis on post-treatment tumors demonstrated a reduction in tumor immunogenicity with a two-fold TMB reduction, a contraction of tumor neoantigen load (Fig. 1j, Supplementary Table), and a loss of a clonal population (Fig 1k, Methods). These results suggest the presence of multiple tumor intrinsic mechanisms that led to tumor recurrence, including the loss of the CAR-targetable antigen along with compromised endogenous T cell recognition, effector function and immune cell exclusion.

### Evidence of tumor resident T cell activity prior to CAR therapy

To understand whether T cells present in the pre-treatment tumor constituted the potential for tumor reactivity, we focused on single cell analysis of tumor infiltrating lymphocytes (TILs). By re-clustering 1192 TILs (excluding NK cells; Extended Data Fig. 3a), we identified five distinct T cell clusters containing cells from pre- and post- treatment tumors: two mixed CD4+/CD8+ clusters of naïve/memory cells (IL7R, CD44), memory/activated cells (CD44, JUNB), and three CD8+ clusters of activated/exhausted cells (IFNG, PDCD1, HAVCR2, LAG3), exhausted cells (PDCD1, HAVCR2, CTLA4) and anergic cells (CBLB, STAT3) (Fig. 2a, Extended Data Fig. 3b,c). The composition of TILs shifted from a mixture of mostly activated/exhausted effector T cells (Pre: 67.4%, Post 3.5%) to a mixture of mostly naïve/memory T cells (Pre: 25.9%, Post: 95.3%).

**Fig. 2.**
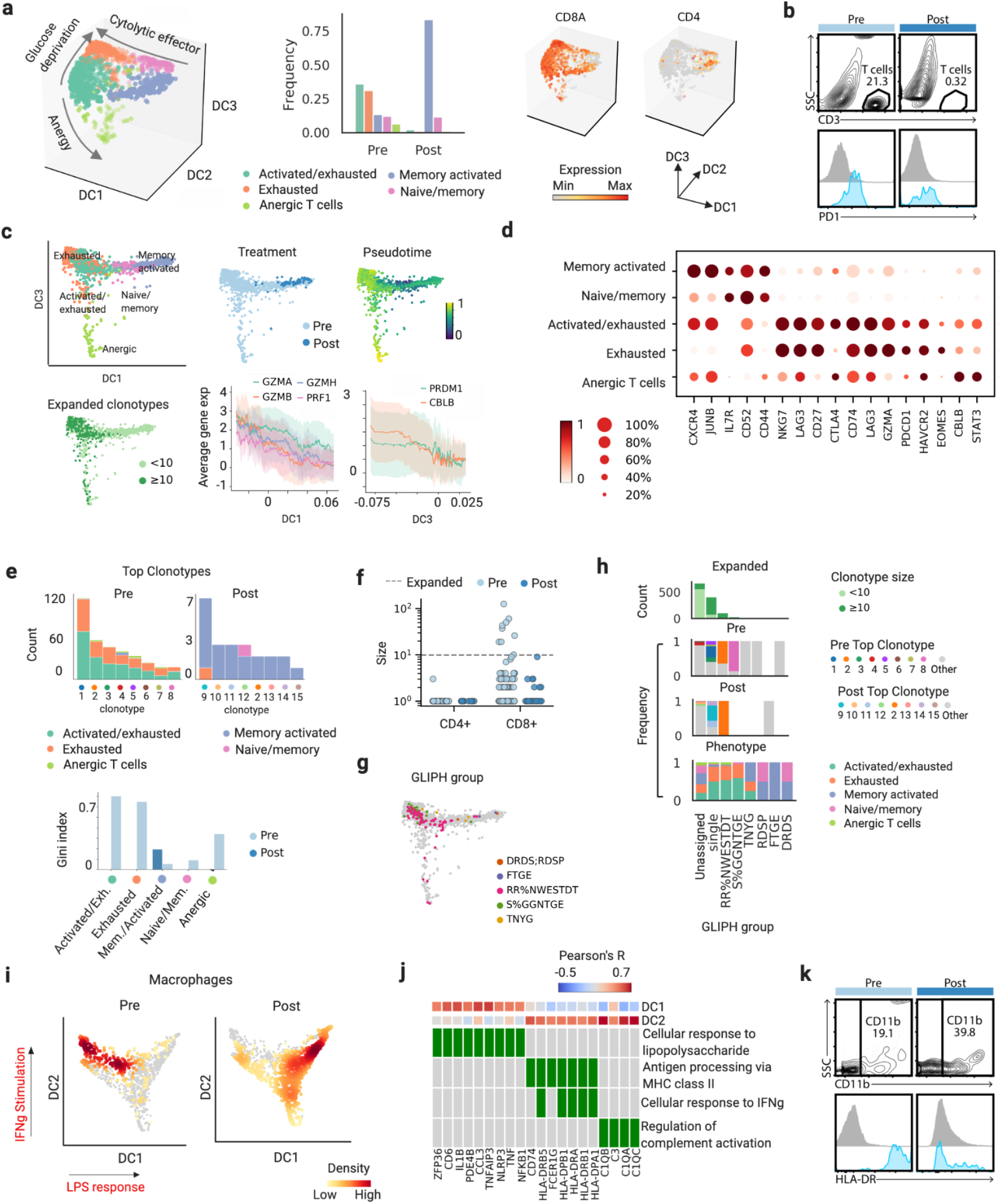
Exhausted CD8+ tumor infiltrating T cells are clonally expanded in the pre-treated GBM tumor. **a**, Three dimensional UMAP of tumor infiltrating T cells present in pre-treatment and recurrent GBM tumors, labelled with inferred cell phenotypes (left) and colored by CD4 and CD8 expression (right). Proportions of each T cell cluster per treatment status (middle). **b**, Flow cytometry characterization of CD3+ and CD3+PD1+ cell loss in post- vs pre-treatment tumors. **c**, Diffusion map of exhausted, exhausted/activated, naïve/memory and memory/activated T cells, using diffusion components 1 and 3, with cells colored by inferred cell type (top), by expanded clonotype (bottom left), by treatment status and pseudotime (top right). Rolling average expression of selected core activation and exhaustion genes as quantified along diffusion components 1 and 3 (bottom right). **d**, Dot plot of genes differentially expressed in T cell clusters, where color represents average expression and size represents percentage of cells expressing. **e,** Distribution of TIL phenotypes for each of the top eight clonotypes in pre- and post-treatment tumors (top). Bar plot of Gini index calculated for each TIL phenotype and treatment status (bottom). **f,** Strip plot of size of CD4+ and CD8+ T cells clones per treatment status. Clones with paired TRA and TRB are assigned CD4+ or CD8+ based on the majority expression of CD4 and CD8A. **g,h**, GLIPH identifies TCR specificity groups defined by local motifs, global similarity, or exact sequence match (single), while some TCRs remain unassigned. **g,** Diffusion map of pre- and post-treatment TILs, with cells colored by TCR specificity groups from GLIPH defined by local or global similarity. **h**, Count bar plots per GLIPH classification colored by expansion defined by clonotype size (≥10) (top). From second to last, frequency bar plots per GLIPH classification colored by unique TRB sequence, top clonotypes as in **e**, and TIL phenotypes. **i**, Diffusion map of tumor resident macrophage population using first two diffusion components. Cells are colored by Gaussian kernel estimate density per treatment status on diffusion map. **j**, Gene Ontology enrichment of highly correlated genes along first two diffusion components for the macrophage population. **k**, Flow cytometry characterization of CD11b+ and CD11b+ HLA-DR+ cell loss in recurrent vs pre-treatment GBM tumors.

We applied diffusion maps (Haghverdi, Buettner, and Theis 2015) to visualize phenotype and order cells in pseudotime: the first diffusion component (DC) separated activated/exhausted cells and memory activated cells and was correlated with T cell cytotoxicity genes (GZMA, GZMB, PRF1; Adjusted p-value=3.0 × 10-56, 6.8 × 10-113, 4.6 × 10-58), while the third DC separated exhausted and activated/exhausted cells from anergic cells and differentially expressed genes associated T cell immunological synapse impairment (Jeon et al. 2004) (CBLB; Adjusted p-value=1.4 × 10-25) (Fig 2c, d). Pre-treatment TILs with high expression of T cell exhaustion markers (PDCD1, LAG3, HVAC) and activation markers (IFNG, JUNB) were associated with CD8+ activated/exhausted cells and suggest the potential for endogenous T cell tumor reactivity (Fig. 2d, Extended Data Fig. 3b,c).

To determine whether pre-treatment TILs had the potential for a productive, tumor specific immune response, we explored the link between T cell clonal expansion, phenotype, and tumor specificity. We measured clonality using the Gini index from single cell TCR sequencing data (Methods) and observed higher clonal expansion in activated/exhausted and exhausted CD8+ T cells compared to all other T cells (Fig. 2e). We identified 9 expanded CD8+ clonotypes (≥10 cells) present in the pre-treatment tumor (Fig 2f). We numbered the top eight clonotypes in pre- and post-treatment tumor as clonotypes 1-8 and clonotypes 9-15 respectively, with clonotype 2 amongst the top clonotypes in both tumors. This shared clonotype encoded for T cells that transitioned phenotypes over treatment course — from activated/exhausted, exhausted or anergic phenotypes to a memory activated phenotype at time of relapse. Although only single-positive (CD39-CD103+) and not double-positive (CD39+CD103+) for tumor-reactive markers (Duhen et al. 2018), the highly expanded pre-treatment top clonotypes exhibit a CD8+ activated/exhausted phenotype. Notably, the proliferation of most expanded clonotype 1 was enabled by expression of TCF-1, a marker associated with progenitor exhausted CD8 T cells with self-renewal potential (Chen et al. 2019; Siddiqui et al. 2019; Beltra et al. 2020; Im et al. 2016)(Extended Data Fig. 3f).

We next used GLIPH (grouping of lymphocyte interactions by paratope hotspots) to identify TIL TCR groups most likely to share antigen specificity (Huang et al. 2020). We found two GLIPH specificity groups with expanded clonotypes (size≥10), including one GLIPH group (RR%NWESTDT) consisting of three unique clonotypes that included clonotype 2 (Extended Data Table 1); cross reference with public TRB sequences indicated that no clonotype had viral specificity (Fig. 2g, h, Extended data Table 2). Similar to recent studies (Yost et al. 2019), we observed that clonally expanded TILs and T cells within a GLIPH group are likely to share a common phenotype, in this case activated/exhausted. Pre-treatment TILs group with respect to antigen-specificity and are clonally expanded prior to CAR T cell therapy. We therefore hypothesize the existence of a pre-existing, antigen specific, though ineffective T cell immune response.

To further characterize tumor immune composition, we re-clustered 1457 tumor associated macrophages (TAMs) and applied diffusion maps (Fig. 2i); the first two diffusion components showed enrichment by treatment status of either upregulation of MHC Class II antigen processing (DC1; Pre), or lipopolysaccharide (LPS) stimulation pathways (DC2; Post) (Fig. 2j). Upregulation of HLA-DR expression (Fig. 2k) and stimulation of MHC II in pre-treatment TAMs, combined with the presence of clonally expanded T cells highlight a coordinated cellular immune response in the pre-treatment tumor, transactivated through reciprocal production of IFNγ.

Combined single cell analysis of malignant and immune cell populations demonstrated that this patient’s pre-treatment tumor — in contrast with the relapsed tumor — was highly immunogenic and infiltrated with clonally expanded, cytotoxic CD8+ T cells that showed evidence of tumor specificity. Notably, a sustained regression of progressing lesions located in the brain and spine (T4 Left temporal, superior gyrus; T5 Left temporal, middle gyrus; T6 Right frontal lobe; T7 Olfactory groove; T8 Lumbar Spine) that occurred following intraventricular infusions of IL13Rα2-CAR T cells, suggests that the therapeutic effect was likely CAR T cell mediated. Since we were unable to biopsy and perform similar immunogenomics analyses on these lesions, our objective turned to characterizing this patient’s T cell mediated immune response in peripheral compartments. We hypothesized that tracing T cell dynamics in the CSF and PB during the treatment course could elucidate the CAR and endogenous T cell contributions towards tumor targeting and sustained clinical response.

### Tracing dynamics in the CSF and PB reveal high frequency T cell dynamics

To track and characterize T cell dynamics in peripheral compartments, we performed bulk T cell receptor beta chain (TRB) sequencing on the CAR T cell product, along with blood and cerebrospinal fluid samples collected longitudinally during treatment (21 timepoints over 228 days, 1057419 TRB sequences) (Fig. 3a,b, Extended data Fig. 4a). We analyzed high-abundance TRBs (Extended Data Fig. 4b-d, Methods), which are less likely to be artifacts of undersampling and to address the difference in signal to noise ratio between the different compartments (Fig 3c). The application of shape-based clustering on high frequency T cell receptor signals revealed eight dynamics classes that emerged in the PB and in the CSF (Fig. 3d,e, Extended Data Fig. 5a,b, Methods). High frequency T cell clones in the CSF clustered into four distinct patterns; peak dynamics (CSF:1; 1513 TRBs) consisting of transient, high frequency T cell clones present at a single timepoint; persistent dynamics (CSF:2; 37 TRBs) include T cell clones detected at multiple time points over the course of therapy; contracting dynamics (CSF:3; 1178 TRB) that decreased in frequency over the treatment course and reactivated dynamics (CSF:4; 134 TRB), whose decrease and expansion coincided with tumor regression and relapse (Fig. 3d, Extended Data Fig. 4e,f). In the PB, where the signal to noise ratio was lower than in the CSF, T cell clonal dynamics exhibited four patterns; one included persistent high frequency dynamics (PB:2 135 TRB), peak dynamics (PB:1, 7 TRB), and T cell clones that expanded and then contracted during recurrence (PB:4; 1 TRB) or expanded in recurrence (PB:3; 9 TRB) (Fig. 3e, Extended Data Fig. 4e,f).

**Fig. 3.**
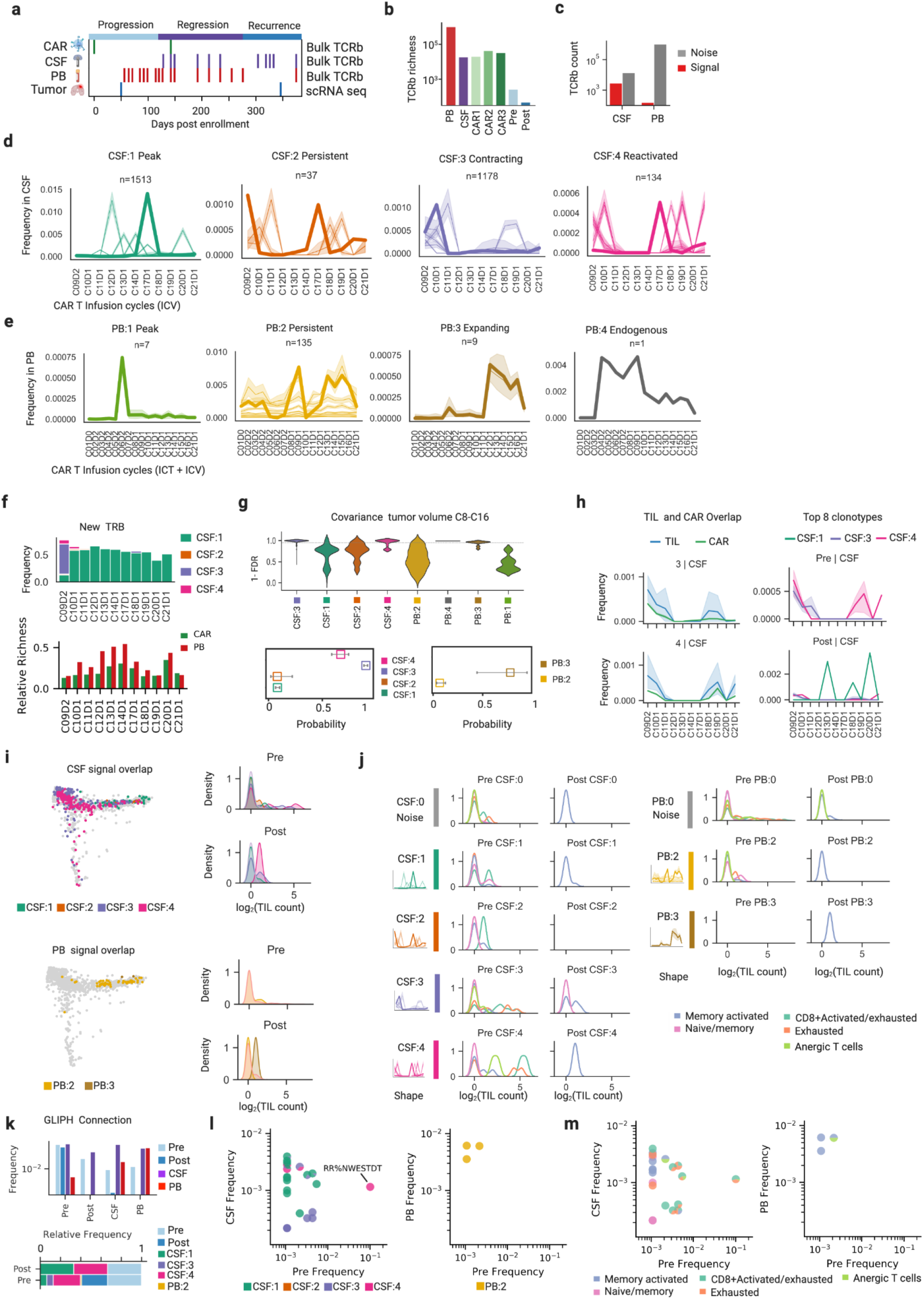
Tracing clonality of tumor resident T cells longitudinally through peripheral compartments uncovers distinc anti-tumor T cell dynamics in the CSF. **a**, Samples collected for bulk TCR beta sequencing over the course of CAR T cell therapy. **b**, Richness of TRB in combined sets of PB, CSF, CAR T product, and TILs. **c**, Size of TRB low frequency (noise) and high frequency (signal) dynamics clusters in PB and CSF. **d**, Mean traces of k-means clusters (with 95 CI band) and k-shape clusters (bold) for CSF dynamics. **e,** Mean traces of k-means clusters (with 95 CI band) and k-shape clusters (bold) for PB dynamics. **f,** Frequency of newly introduced TRB sequences into the CSF by each signal dynamics class (Methods) (top). Number of TRB sequences newly introduced by peak dynamics CSF:1 that overlap with CAR T product sequences (CAR 1, cycles 1-11, CAR 2/3: cycles 12-21) and closest previous peripheral blood timepoint. (Methods, Extended Data) (bottom). **g**, p-values, adjusted for FDR by the Benjamini-Hochberg procedure, for significance of rejection of independence test between PB and CSF signal dynamics and tumor volume during tumor regression phase (left). Estimate for the probability that a random member of each signal class is not independent from the tumor volume, as determined by the calculated p-values and FDR of 0.5, with error bars showing confidence intervals calculated using the Agresti-Coull method (right). **h**, TRB traces of TRB sequences that overlap with TIL and CAR T product sequences, in CSF dynamics classes 3 and 4 (left). Traces of TRB sequences corresponding to top 8 clonotypes and CSF dynamics classes 1, 3, and 4 (right). **i,** Time series of top clonotypes from pre- and post-treatment tumors that overlap with CSF and PB noise and signal dynamics.

### Intraventricular CAR T infusions introduce new T cell diversity into the CSF

We further characterized high frequency T cell dynamics in the CSF to examine whether locoregional delivery of CAR T cells could modulate T cell immunity in the CNS. We calculated the frequency of new T cell receptor clones over the treatment course and found that peak dynamics, in contrast to other high frequency CSF dynamics, introduced new T cell receptor diversity into the CSF (Fig. 3f). Namely, the peak dynamics’ T cell receptor repertoire was primarily composed of TRBs that overlapped with the PB following every infusion (24% CAR and PB, 39% PB only, 2.4% CAR only), suggesting migration from the PB into the CSF (Fig. 3f, Extended Data Fig. 6f, Methods).

Previous results have highlighted that intraventricular CAR T cell infusions drive the amplification of cytokines related to T cell activity and chemoattraction in the CSF (Brown et al. 2016). We hypothesize the peak dynamics represent CAR T cell infusions and new T cell recruitment from the PB—and highlight CAR T cell potential to transiently modulate endogenous T cell activity in the CSF. Compared to peak dynamics, T cell clones in the persistent, contracting and reactivated dynamics matched more with TRB sequences from either CAR T infusion products or tumor resident T cells (CSF:1 27%, CSF:2 62%, CSF:3 31%, CSF:4 51%) (Extended Data Fig. 6g). We conjecture the locoregional expansion and contraction of these T cell clones in the CSF associate with tumor antigen exposure and killing. Despite sustained intraventricular infusions of CAR T cells, richness of the CSF TRB repertoire decreased during tumor regression, suggesting the existence of a tumor-targeting T cell dynamics class to explain this loss of T cell diversity (Extended Data Fig. 6ab).

### T cell clonal dynamics in the CSF covary with tumor regression

To evaluate tumor killing potential of locally expanded T cells, we sought to identify whether high frequency T cell clones in PB or CSF tracked with tumor volume regression as an indicator of T cell mediated immunity targeted toward regressed tumors. We performed covariance independence testing of each T cell clone timeseries with respect to tumor volume during regression (cycles 8-16) and found that CSF contracting and reactivated T cell dynamics covaried with tumor volume with high probability (CSF:3 p=0.91, CI=−0.02/+0.02; CSF:4 p=0.67, CI = −0.08/+0.07) (Fig. 3g, Methods). Strikingly, five of the top 8 expanded pre-treatment clonotypes traced with contracting and reactivated dynamics (CSF:3 clonotypes 3,6,7; CSF:4 clonotypes 2,4) (Fig. 3h,i).

### Tracing pre- and post-treatment TILs in peripheral compartments

To explore the extent to which pre-treatment tumor resident T cells could constitute the basis for a secondary T cell response targeting regressed tumors, we traced T cell clones from pre- to post-treatment tumors and in peripheral (PB) and central (CSF) immune compartments by matching clonotypes based on TRB sequences. We calculated signal to noise ratio of matched TRBs in tumor and peripheral compartments and found that a large proportion of TILs were maintained at low frequency in blood compared to the CSF (CSF:S/CSF:N = 1.05; PB:S/PB:N = 0.147) (Fig. 4a,b, Extended data Fig. 6d). Most of the clonally expanded TILs (11 out of 15 top clonotypes)—with the exception of clonotype 13 that expanded in PB during tumor regression and trafficked to the post-treatment tumor— overlapped with PB noise dynamics (Fig 3i). Conversely, in the CSF, these T cell clones—matched with CD8+ activated/exhausted, exhausted and anergic tumor resident phenotypes—and tracked with high frequency contracting or reactivated dynamics (Pre TIL: 5/8; Post TIL: 3/8 top 8 clonotypes) (Fig. 4a,c, Extended Data Fig. 6d). The most clonally expanded pre-treatment clonotype (clonotype 1), in contrast, appeared in the PB at low frequencies and was absent in the CSF, suggesting this clonotype might not have been selected for due to lack of antigenic specificity in the case of a secondary immune response.

**Fig. 4.**
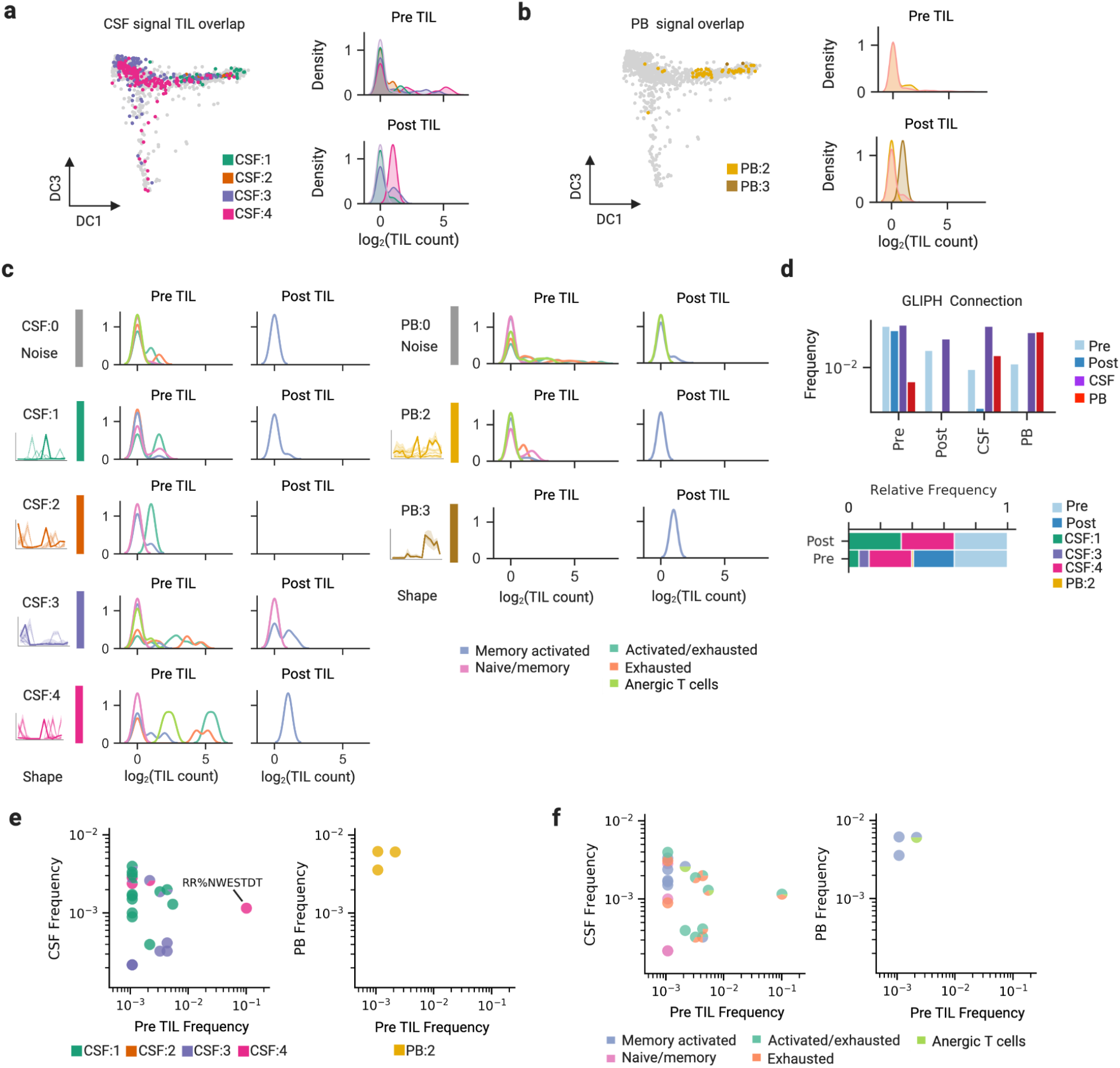
Reactivated and contracting T cell dynamics in CSF overlap with antigen specific, exhausted and expanded pre-treatment TILs. **a,b,** UMAP of TILs colored by TCR shared with each dynamics class (left) for CSF in **a** and PB in **b**. Corresponding kernel density approximation of overlapping Pre and Post TIL count per signal dynamics (right). **c**, Kernel density approximation of Pre and Post TIL count per phenotype, grouped by overlap with TCRs from CSF (left) or PB (right) noise and signal dynamics. **d**, Bar plot showing the aggregated frequency of TCR connecting tumor, CSF, and PB compartment via GLIPH groups with local or global similarity (Methods) (top). Bar plot showing relative frequency of TRB within pre- and post-treatment tumor connected to tumor, CSF dynamics, or PB dynamics via local or global similarity (bottom). **e,f,** Scatter plot of GLIPH classes connecting the pre-treatment tumor to CSF (left) or PB (right). Frequencies of TCRs are summed per GLIPH class, and colored by distribution of dynamics classes in **e**, or by frequency distribution of TIL phenotype in **f**.

To characterize antigen specificity shared between TILs and expanded circulating T cells in CSF and PB, we calculated the intra and inter-compartment connectivity of GLIPH-identified TCR antigen specificity groups (Fig. 4d, Methods). Our analysis showed greater antigen specific connections between pre- and post-treatment TIL and CSF high frequency clones —highlighting that shared antigen specificity is more directly observed between tumor TILs and CSF compared to PB. To compare clonal expansion and dynamics of GLIPH groups in the periphery, we aggregated the cumulative frequencies for each GLIPH connection between tumor and CSF or PB. T cell clones in GLIPH specificity group RR%NWESTDT traced through CSF reactivated dynamics (CSF:4) — contracting with spinal tumor regression and re-expanding with relapse tumor growth (Fig. 4e). Notably, it included a shared, clonally expanded clonotype (2), present in both pre and post treatment tumors. Investigation of T cell phenotypes per GLIPH group showed that most CSF GLIPH groups expanded at the tumor exhibited mixed activated/exhausted phenotype, while PB GLIPH groups exhibit memory phenotype with no expansion (Fig. 4f). Together, these results suggest the existence of shared antigens and the potential to observe the dynamics of immunological memory in the CSF.

Tracing clonality and antigen specificity of tumor resident T cells in peripheral compartments highlight differences in anti-tumor dynamics in peripheral and central adaptive immune compartments. These results suggest that the peripheral blood represents persistent maintenance of clonally expanded, cytotoxic TILs, similar to previous studies (Chu et al. 2019). By contrast, the CSF reveals local T cell mediated immunity with potential cytotoxicity towards shared antigens in the (resected) pre-treatment tumor and responsive tumors. While this phenomenon of higher T cell clone sharing between brain lesions and CSF than PB was observed in multiple sclerosis (Salou et al. 2015) an antigen-driven autoimmune neurological disease, this is the first tracking of lesion repertoire comparing peripheral and central TCR repertoire for glioblastoma.

## DISCUSSION

Here, we performed combined single cell and TCR clonotype analysis on GBM tumors, cerebrospinal fluid and peripheral blood samples collected longitudinally from a patient with multifocal glioblastoma treated with IL13Rα2-targeting CAR T cells that exhibited a complete response to therapy and then subsequently recurred at new sites. This patient’s disease course provided the opportunity to uncover dynamics of T cell mediated immunity during response and then subsequent recurrence. Our analysis revealed hallmarks of an inflamed tumor immune microenvironment pre-CAR T cell therapy — including the presence of CD8+ cytotoxic, clonally expanded and antigen-specific T cells that tracked in the cerebrospinal fluid. These expanded then subsequently contracted with tumor volume, or re-expanded upon tumor recurrence, suggesting the existence of a T cell immune response that combined with CAR T cells to target unresected and progressing spinal and brain tumors. Although regressing tumors responding to CAR T cell therapy could not be queried to elucidate changes in the TME associated with response, the evaluation of post-CAR recurrent tumors provides insights into tumor features that are less responsive to CAR T cell therapy. Our demonstration that a productive CAR T cell response, as achieved in this unique responder, is associated with the induction of endogenous immune reactivity provides a paradigm for improving CAR T cells activity against solid tumors generally. Other examples of T cell therapy modifying host immune responses to achieve dramatic clinical responses have been observed (Hunder et al. 2008; Hegde et al. 2020), including a recent report demonstrating alterations in T cell repertoires and antibody reactivity following HER2-CAR T cell therapy that was associated with durable remission of a patient with rhabdomyosarcoma (Hegde et al. 2020). Our findings also suggest that an inflamed tumor microenvironment may be more responsive to CAR T cell therapy, as has been shown for checkpoint immunotherapy (Ayers et al. 2017), a notion that requires further clinical confirmation.

In this study, we demonstrate that liquid biopsy of the cerebrospinal fluid provides a safe and effective strategy for real-time monitoring of immune dynamics in the central nervous system (Miller et al. 2019). T cell dynamics associated with intraventricular infusions of CAR T cells and tumor reactive T cells were more distinguishable in the CSF in contrast to the PB. We propose that the CSF is an informative liquid biopsy, and could be used in combination with methods that investigate TCR-antigen specificity (Scheper et al. 2019; Gee et al. 2018) to capture and understand locoregional T cell dynamics that are more closely related to tumor microenvironment in patients with CNS disease. As observed in the recurrent tumor, the lack of T cell activity and antigen specificity presents a problem in the field of immunotherapy; however, monitoring the evolution of antigen specificity via the CSF may guide the optimization of combinatory targeted immuno- or chemotherapy. Increased cytokine activity, T cell recruitment, and anti-tumor efficacy during intraventricular delivery emphasized the therapeutic potential to manipulate CNS immunity via the CSF, prompting future investigation of its effect on T cell priming in the CNS meningeal lymphatic system (Louveau et al. 2015, 2018) and potential synergy with lymphoangiogenic enhancement to increase T cell priming (Song et al. 2020).

## METHODS

### Human participants

Tumor, CSF and blood samples were collected from a patient participating on our IL13Ra2-CAR T cell phase I clinical trial (NCT02208362 and single subject protocol), with treatment history and clinical outcome previously reported1. This clinical study was approved by the City of Hope Institutional Review Board and the research participate gave written informed consent.

### Tumor dissociation

Leftover fresh tumor specimens were obtained at the time of tumor resections as necessitated by clinical need. Tumors were dissociated by mincing and digesting with the Tumor Dissociation kit (Miltenyi Biotec) per the manufacturer’s instructions using a gentleMACS dissociator (program h_tumor_01 and h_tumor_2). DMEM F/12 supplemented with B-27serum and heparin was used in place of RPMI. The cell suspension was then passed through a 40-μm filter and pelleted by centrifugation at 300g for 10 min. Cells were then resuspended in Crystor10 (Biolife Solutions, Inc) and frozen in 0.5ml per cryovial and subsequently stored in liquid nitrogen freezer until further processing. Freshly dissociated tumor cells were then thawed and processed for scRNA-seq.

### Flow cytometry and immunohistochemistry

Slides are first deparaffinized on the Ventana Discovery Ultra. Once the paraffin is removed from the slides and tissue, the tissue gets treated with antigen retrieval and heat to open the binding sites for linking. Once the binding sights are available the tissue is incubated with the first primary antibody. Next, the linking secondary antibody is applied to the slides to link the primary antibody to the chromogen. Now the chain can be completed with chromogen, which in this case is CD66B in DAB, CD8 in Purple, CD4 in Teal, CD68 in RED, and GFAP in Yellow. Lastly the slides are counter stained with Hematoxylin to show the surrounding cell morphology. These steps are all completed on a Ventana Discovery Ultra staining machine. The last step is to coverslip each slide for tissue longevity and increased visualization under a bright field microscope. Each antibody is stained to completion before the next antibody is started. For IL13RA2 (1:600, R&D Systems AF146), HER2 (1:200, Dako A0485), and EGFR (1:100, Invitrogen 28-0005) staining, formalin-fixed, paraffin-embedded tissues were cut at 5uM and stained using Envision+System-HRP (DAB) chemistry and AutostainerPlus hardware (both Dako) by the City of Hope Pathology Core shared resource. Cells were stained with fluorochrome-conjugated antibodies specific for human CD3, CD45, CD11b, HLA-DR (BD Biosciences), PD-1 (BioLegend), and with DAPI (Invitrogen) used as a viability dye. Samples were then run on a MACSQuant Analyzer (Miltenyi Biotec Inc.) and analyzed using FlowJo (v. 10.7) (FlowJo, LLC).

### Tumor volume and interpolation

The last dynamic phase of the dynamic contrast enhanced (DCE) sequence was used to identify lesions. Regions of interest (ROIs) were drawn manually on each slice where the lesion was visible in the transaxial plane, covering the enhancing and non-enhancing components of each lesion. Regions of normal vasculature were avoided whenever possible. The sum total number of voxels in the ROI was used as a measure of total contrast enhancement (CE) and of total tumor volume. Only the voxels inside the manual ROI that showed a confidence level of at least 80% in the DCE kinetic model fit were included in the final tumor volume (Sahoo et al. 2013). This approach includes both early- and late-phase enhancing voxels and excludes voxels with low or noisy enhancement. To interpolate tumor volume between observed timepoints, we used linear interpolation implemented in SciPy (v.1.2.3) (Virtanen et al. 2020).

### Bulk RNA sequencing

RNAs were converted to cDNA libraries using Ribo-ZeroTM rRNA Removal Kits (Illumina) and TruSeq RNA Sample Preparation Kit V2 (Illumina) following manufacturer’s recommendations. Libraries were sequenced on the Illumina HiSeq 2500 with paired-end 101 bp read mode at the City of Hope Integrative Genomics Core facility.

### Exome sequencing

Exome capture was performed and sequenced on an Illumina HiSeq 2500 with paired-end 100-base pair (bp) sequencing, with an average 100-fold coverage. Illumina adapters were added to genomic DNA to make a library for paired-end sequencing (Illumina Inc., San Diego, CA). Fragments with approximately 200–250 bp insert DNA were selected and amplified. The library was hybridized to biotinylated cRNA oligonucleotide baits from the SureSelect Human All Exon kit V6+somatic probe sets (Agilent Technologies Inc., Santa Clara, CA), and amplified for 12 cycles. After purification, the library was paired-end (100+100bp) sequenced using an Illumina Hiseq 2500 (Illumina Inc., San Diego, CA).

### HLA Typing

High-resolution HLA typing was performed by City of Hope’s HLA Laboratory using a combination of the following methods (resolving common ambiguous genotypes): sequencing-based typing, PCR- SSOP (sequence-specific oligonucleotide probes), and/or PCR-SSP (sequence-specific primers). Different alleles from the same G groups were considered matched.

### Bulk TCR sequencing

Survey level sequencing of the TRB gene was performed using the immunoSEQ platform (Adaptive Biotechnologies) on genomic DNA extracted from peripheral blood, cerebrospinal fluid and CAR T products. Genomic DNA input amount ranged from 395.702 ng to 2039.886 ng in peripheral blood samples, 99.816 ng to 593.866 ng in the cerebrospinal fluid, and 399.510 ng to 400.466 ng in CAR T products (Extended data, Table). Raw data was exported from the immunoSEQ Analyzer; only data from productive TCR rearrangements was considered in downstream analysis. On average, 110943 TCR templates were detected from peripheral blood samples (range 46139–255259), representing an average of 84962 unique clonotypes (range 36396–203184). For cerebrospinal fluid samples, on average 2218 TCR templates were detected (range 123–9234), representing an average of 1800 unique clonotypes (range 116– 6736). For CAR T products, on average 73207 templates were detected (range 65678–78528), representing an average of 36051 unique clonotypes (range 23654–48816).

### Preparation of scRNA-seq libraries

The single cell RNA-seq and TCR-seq libraries were prepared using the 10x Single Cell Immune Profiling Solution Kit, according to the manufacturer’s instructions. Cells were washed once with PBS containing 0.04% weight/volume bovine serum albumin (BSA) (InvitrogenTM UltraPure BSA, Cat. No. 2616) and resuspended in PBS to a final concentration of 100–1000 cells per μl as determined by hemacytometer. Cells were captured in droplets at a targeted cell recovery of 10,000 cells. Following reverse transcription and cell barcoding in droplets, emulsions were broken and cDNA purified using Dynabeads MyOne SILANE followed by PCR amplification (98 °C for 45 s; 13–18 cycles of 98 °C for 20 s, 67 °C for 30 s, 72 °C for 1 min; 72 °C for 1 min). Amplified cDNA was then used for both 5′ gene expression library construction and TCR enrichment. For gene expression library construction, 40 ng of amplified cDNA was fragmented and end-repaired, double-sided size-selected with SPRIselect beads, PCR-amplified with sample indexing primers (98 °C for 45 s; 14–16 cycles of 98 °C for 20 s, 54 °C for 30 s, 72 °C for 20 s; 72 °C for 1 min), and double-sided size-selected with SPRIselect beads. For TCR library construction, TCR transcripts were enriched from 2 μl of amplified cDNA by PCR (primer sets 1 and 2: 98 °C for 45 s; 10 cycles of 98 °C for 20 s, 67 °C for 30 s, 72 °C for 1 min; 72 °C for 1 min). Following TCR enrichment, 5–50 ng of enriched PCR product was fragmented and end-repaired, size-selected with SPRIselect beads, PCR-amplified with sample-indexing primers (98 °C for 45 s; 9 cycles of 98 °C for 20 s, 54 °C for 30 s, 72 °C for 20 s; 72 °C for 1 min), and size-selected with SPRI select beads.

### Single cell sequencing

The single cell RNA libraries were sequenced on an Illumina NovaSeq to a minimum sequencing depth of 25,000 reads per cell using read lengths of 26 bp read 1, 8 bp i7 index, 98 bp read 2. The single-cell TCR libraries were sequenced on an Illumina MiSeq or NovaSeq to a minimum sequencing depth of 5,000 reads per cell using read lengths of 150 bp read 1, 8 bp i7 index, 150 bp read 2.

### Bulk RNA-seq data processing

Using cutadapt (version 1.9.1) (Martin 2011), raw RNA-Seq reads were quality trimmed (cutoff=19, 5’ and 3’, single-end mode) before adapters trimming in pair-end mode. Fusions were detected per sample using two steps: Potential fusion junctions were found during kallisto (version 0.46.0) pseudoalignment to GRCh38 cDNA (Ensembl release 96) using fusion mode (Bray et al. 2016), then fusion junctions were filtered by pizzly (https://github.com/pmelsted/pizzly) based on Gencode genome annotation (version 30) to generate a FASTA file of fusion transcripts (Melsted et al., n.d.). After filtering redundant fusions, the combined set of fusion transcripts from multiple samples were added to GRCh38 cDNA to build a new kallisto index, used for kallisto re-quantification of transcript and fusion abundance with 100 bootstraps per sample. Our fusion quantification workflow available at https://github.com/vdjonsson/rna-seq-pizzly was written with Snakemake (Köster and Rahmann 2018). Normalization and differential analysis were performed with sleuth (version 0.30.0) using gene-level p-value aggregation of likelihood ratio test comparing a full model containing treatment label and a reduced model ignoring treatment label (Pimentel et al. 2017; Yi et al. 2018). For downstream analysis, the transcript count matrix, in Transcripts per Kilobase Million (TPM), was summed per gene and transformed with log_2_(*TPM*+0.5). Log-transformed gene-level values were averaged per condition before taking the difference (Pre - Post) to calculate log_2_(FC). Significantly up- or down-regulated genes with *qval* < 0.05 and |log_2_(FC)| > 1 were enriched for Gene Ontology Biological Processes using GSEApy (version 0.9.6) (Fang n.d.).

### Data processing of exome libraries

The paired-end sequences from tumor and normal samples were aligned to GRCh38 (hg38) using Novoalign (http://www.novocraft.com) with default settings. Only reads aligned to a unique genomic location were kept for further analysis. Aligned reads were piled up using Samtools version 0.1.19. Variants (SNPs and short indels) were identified using MuTect2 v2 and further annotated with the GATK. Variant functional annotation was performed using AnnoVAR. Germline copy number variants (CNV) were identified using CoNifer, and virtual somatic copy number variants were identified using Bioconductor package DNAcopy. Neoepitopes were identified using pVAC-seq and a peptide–MHC pair was considered a neoepitope if the peptide was found to bind to the MHC allele with less than 500 nM binding strength and the wild-type cognate peptide bound to the same allele with a binding strength greater than that of neoepitope. Tumor clonal analysis was performed using lumosVar v2 (Halperin et al. 2019).

### Preprocessing of scRNA-seq data

The cellranger count (10x Genomics, version 3.0.1) was used to align scRNA-seq reads to GRCh38 reference genome, generate gene count matrix, and filter barcodes. We used filtered gene-barcode matrices containing only cell barcodes determined by cellranger’s cell- calling algorithm that utilizes total UMI threshold and RNA profile model. Pre1 had a median of 1983 genes, 5092 unique transcripts per cell from a total of 3526 cells. Post2 had a median of 1463 genes, 2987 unique transcripts per cell from a total of 7116 cells.

### Processing of single cell count matrix

Single cell data processing was performed with Scanpy (version 1.4.4) (Wolf, Angerer, and Theis 2018). We removed genes with zero expression, miRNA genes with “MIR” prefix, and genes annotated with lincRNA or antisense gene biotypes to give a final set of 17434 genes. Gene matrix was total-count normalized to 10000 reads per cell and then log transformed. We removed cells with less than 300 genes, less than 1000 total UMI counts, greater than 20% mitochondrial RNA, or two standard deviation below average complexity (ratio of UMI count to gene count). After filtering, 8152 out of 10642 cells remained in the combined dataset (76.6%). With score_genes function, each cell was assigned a heat-shock score calculated using genes annotated with Gene Ontology biological processes term ‘response to heat’. Using score_cell_cycle function, we assigned each cell a cell cycle phase based on S and G2M scores calculated with previously published gene set (Tirosh et al. 2016). To remove batch effected and other unwanted biological variation, we regressed out total UMI count, percent mitochondrial RNA, S score, G2/M score, and heat-shock score from the log-normalized matrix. Doublets identified per sample using Scrublet40 were excluded from downstream analysis of cell subpopulations. Additionally, we trained a linear-decoded variational auto-encoded model (scVI version 0.4.0) on raw counts to obtain normalized, batch corrected, imputed data (Lopez et al. 2018). For visualization, we used the resulting mean of the negative binomial distribution following regressing out of unwanted variation (cell cycle, % mitochondrial RNA, heat-shock).

### Dimension reduction and clustering

For the whole or subset (Malignant, T cell, Monocyte) data, we performed the following dimension reduction procedures. The regressed log-normalized data is decomposed by principal component analysis. The number of principal components (PCs) to retain was selected using knee-detection in kneed. These selected PCs were used as input to compute a neighborhood graph of k=15 nearest neighborhoods (method=umap). Leiden clustering algorithm was applied to the neighborhood graph over a range of resolution (0.1-1.2) and the optimal clustering resolution was selected for a high mean silhouette score. Using published gene signatures for glioblastoma and general immune populations (Azizi et al. 2018; Darmanis et al. 2017), clusters were annotated based on average detection rate (non-zero expression normalized by number of genes). Differential gene expression analyses between clusters or other observational groups were performed on non-regressed log- normalized data using t-test with Benjamin-Hochberg correction. Significantly up- and down-regulated genes (*qval*< 0.005, |log_2_(FC)| > 1) were enriched for Gene Ontology Biological Processes using GSEApy (version 0.9.6).

### Diffusion component analysis

We recomputed the neighborhood graph of k=15 nearest neighborhoods using a Gaussian kernel on the basis of the selected PCs. The neighborhood graph is used as input to find fifteen nonlinear diffusion components. The 0th component is trivial and discarded. For each component, highly correlated genes with |*R*| > 0.5 were identified and enriched for pathways as described above. For T cell subset, a randomly selected naïve/mem T cell was selected as the root cell for computing diffusion pseudotime using fifteen diffusion components and 1% of total number of cells as minimum group size.

### Single cell CNV inference

Copy number variation (CNV) estimation was performed per sample using inferCNV (https://github.com/broadinstitute/inferCNV) that sorts genes by chromosomal location and applies a moving average (window of 100 genes) to the relative expression values. Raw UMI counts of non-doublet cells were used as input into the algorithm and hierarchically clustered per group after processing. Genes with expression in less than 3 cells or a mean number of 0.1 counts across cells were excluded. Cells in non-malignant clusters (T cell, Monocytes, Oligodendrocytes, Vascular) were used to define a normal baseline karyotype, of which the average CNA profile was subtracted from that of the malignant cells (Neoplastic). During the denoising step, the residual normal signal is removed. Furthermore, a six-state CNV Hidden Markov Model was trained to predict CNV regions per defined cell type (analysis_mode=samples).

### Processing of TCR-seq data

The cellranger vdj (10x Genomics, version 3.0.1) was used to align TCR-seq reads to GRCh38 reference and annotate clonotype consensus. We pooled clonotypes from all samples and renamed the clonotypes defined by unique pairs of alpha chain (TRA) and beta chain (TRB) amino acid sequences. Cells with multiple alpha or beta chains were assigned multiple clonotypes according to all possible pairings of TRA/TRB.

### Glioblastoma cellular states

Malignant cells were scored with function score_gene using six meta-modules (MES1-like, MES2-like, NPC1-like, NPC2-like, AC-like, OPC- like) (Neftel et al. 2019). Malignant cells were assigned to the highest scored meta-modules. We constructed a two-dimensional representation of the four GBM cellular states (OPC, NPC, AC, or MES) as outlined in *Neftel et al.*

### Bulk TRB sequencing analysis

To examine high frequency (signal) dynamics of TRB sequences over the course of therapy, we identified sequences with low frequency (noise) and removed them from the dynamics analysis. For the more discrete CSF dataset, we kept all TRBs with frequency greater than the mode frequency per time point and defined the union set of these TRB sequences as signal. We identified noise time series in the PB dataset as those that have infinity norms below a threshold determined by the minimum of the infinity norm multimodal density function. Following the identification and removal of noise, the resulting signal time series were clustered using k-means scikit-learn (version 0.23) in python. TRB time series in CSF were z-score normalized prior to k-means clustering. Using an elbow plot of the total within cluster sum of squared distances between time series against the number of clusters, the optimal number of clusters was determined for TRB time series in peripheral blood and z-score normalized cerebrospinal fluid separately. The TRB time series in peripheral blood and cerebrospinal fluid were then clustered separately with their respective optimal number of clusters. The mean representative time series for each of the computed k-means clusters were clustered using the k-shape time series clustering algorithm for PB and CSF separately in order to determine dominant dynamics classes of TRBs. All TRB time series analysis was implemented with scikit-learn (version 0.23) and python (version 3.8.3).

### GLIPH analysis

GLIPH analysis (version 2) was performed on the combined dataset of TCR bioidentity (TRB, TRBV, TRBJ, TRA) from CSF signal, PBMC signal, and pre- and post-treatment tumors (Huang et al. 2020). Using a CD4/CD8 TCR reference, GLIPH analysis identified TCR specificity groups based on enriched local motifs or global similarity (all amino acids interchangeable). For longitudinal bulk TCR sequencing input, TRA was omitted and frequency was summed over time. To prevent false positive GLIPH groups resulting from TRA dropouts, single cell TRA were imputed with the most frequently found TRA for the corresponding TRB if detected.

### Combined Statistical analysis

Tumor volume was divided into three dominant timeframes: tumor growth cycles 1-8, tumor regression, cycles 8-16, and tumor relapse cycles 16-21. TRB frequencies were extracted for cycles 8-16 during tumor regression, and tested for covariance with respect to tumor volume using a chi squared independence test and corrected for false discovery rate (FDR) using Benjamini-Hochberg procedure. Probability that a random TRB of each shape class is not independent from the tumor volume was estimated using calculated p-values and FDR threshold of 0.05. Confidence intervals were calculated for these probabilities of each shape class using the Agresti-Coull method.

## Data availability

All ensemble and scRNA-seq data is deposited in the GEO and will be available after IRB approval. Exome-sequencing data are deposited in the Sequence Read Archive (SRA) and will be available upon IRB approval. Bulk TCR-seq data can be accessed through the ImmuneACCESS database of Adaptive Biotechnologies (URL). All other relevant data will be available upon IRB approval. All figures created and assembled using biorender.com.

## Code availability

Code for single cell analysis and combined analysis of T cell receptor sequencing dynamics analysis is available at https://github.com/vdjonsson/scimmunity.

## Acknowledgements

We thank members of the T cell Therapeutics laboratory for discussions; Xiwei Wu, Hanjun Qin, Shu Tao at the City of Hope Bioinformatics Core, at the City of Hope Blood HLA core. We thank Russell Rockne and Prativa Sahoo for tumor volume measurements. This work was supported by the California Institute of Regenerative Medicine (CIRM; CLIN2-10248), Parker Institute for Cancer Immunotherapy, and a sponsored research agreement from Mustang Bio. V.D.J. was supported by an award from the National Cancer Institute of the National Institutes of Health (K12CA001727). D.W. is supported by NCI fellowship 5F99CA234923-02. Sequencing was performed by the City of Hope Bioinformatics Core Facility, Immunohistochemistry was performed by Research Pathology, supported by the National Cancer Institute of the National Institutes of Health under grant number P30CA033572. The content is solely the responsibility of the authors and does not necessarily represent the official views of the National Institutes of Health.

## Authors’ Contributions

V.D.J., S.J.F., B.B. and C.E.B. conceived the project. V.D.J. and C.E.B. led the project. V.D.J. led the bioinformatic, computational and data analytic methodology development and analysis. V.D.J, D.A. and C.E.B led the methodology development and analysis. V.D.J., R.N. and N.D. designed and implemented analytic and computational pipelines. R.N.implemented the single cell RNA sequencing pipeline. J.H. consulted on statistical analysis. R.W., B.A., R.S., L.W., D.A. and D.W. performed sample preparation and wet lab experiments. First manuscript draft: V.D.J. and R.N. Final manuscript: V.D.J., R.N, N.D and C.E.B. with input from all authors.

## Corresponding authors

Correspondence to Vanessa Jonsson; and Christine Brown.

## Ethics declarations

### Competing interests

S.J.F. and C.E.B. receive royalty payments from Mustang Bio; all other authors declare no competing interests.

---

**Extended Data Fig. 1.**
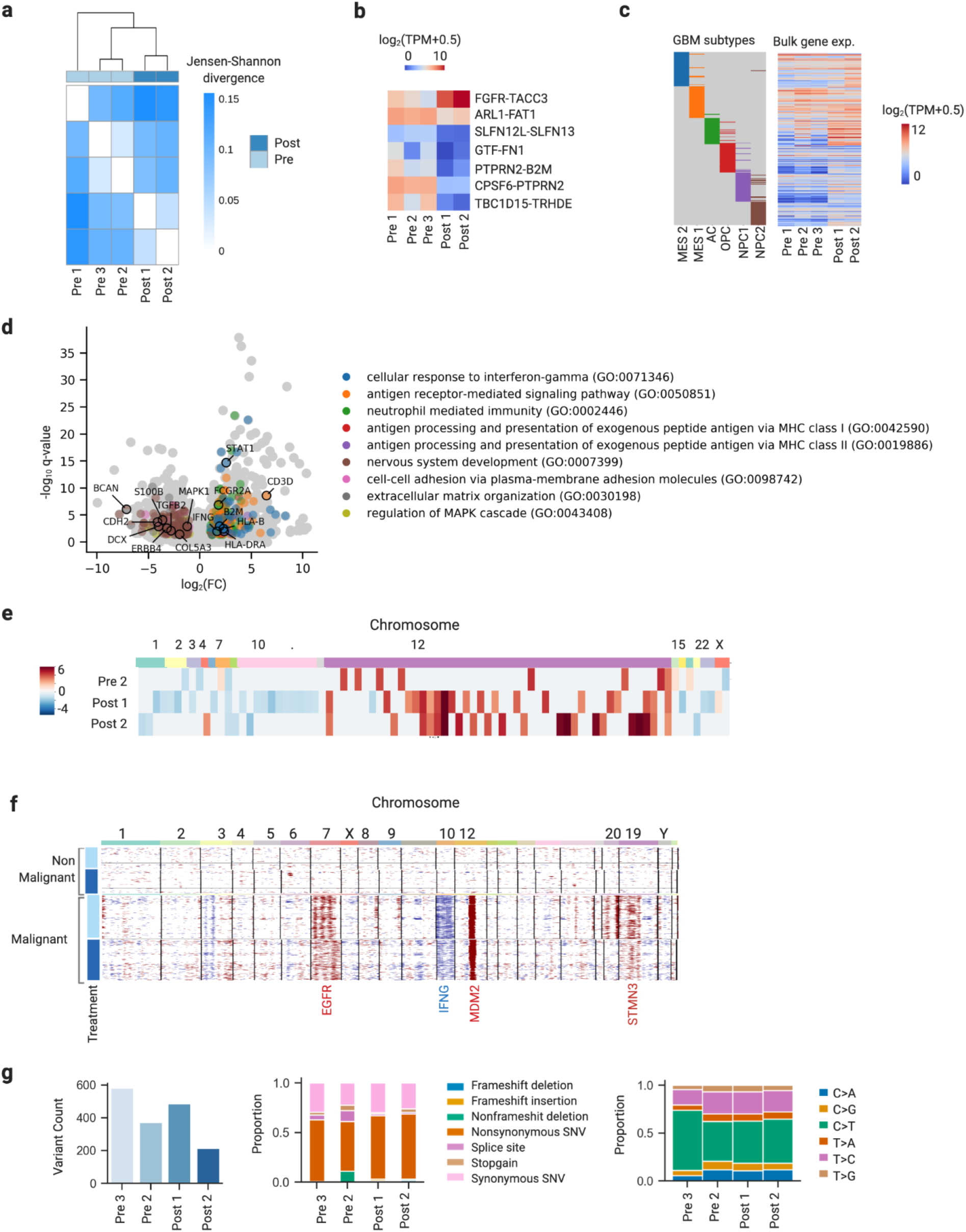
Transcriptomic and genetic landscape of GBM tumors pre- and post-CAR T therapy. **a**, Hierarchical clustering of pre- and post-treatment bulk RNA-seq tumor samples based on similarity measure, calculated as Jensen-Shannon divergence. **b**, Heatmap of top differentially expressed fusions detected in bulk RNA- Seq comparing pre- and post-treatment tumors. **c**, Heatmap of bulk expression of GBM meta-modules. **d**, Volcano plot of differential expression results comparing bulk expression of pre- and post-treatment tumors. Selected GO Biological Processes significantly upregulated in each condition are colored and annotated by representative genes. **e**, Inference of CNV from single cell RNA sequencing data, with rows corresponding to single cells. **f**, Summary of mutational burden, mutation frequencies and mutational signatures.

**Extended Data Fig. 2.**
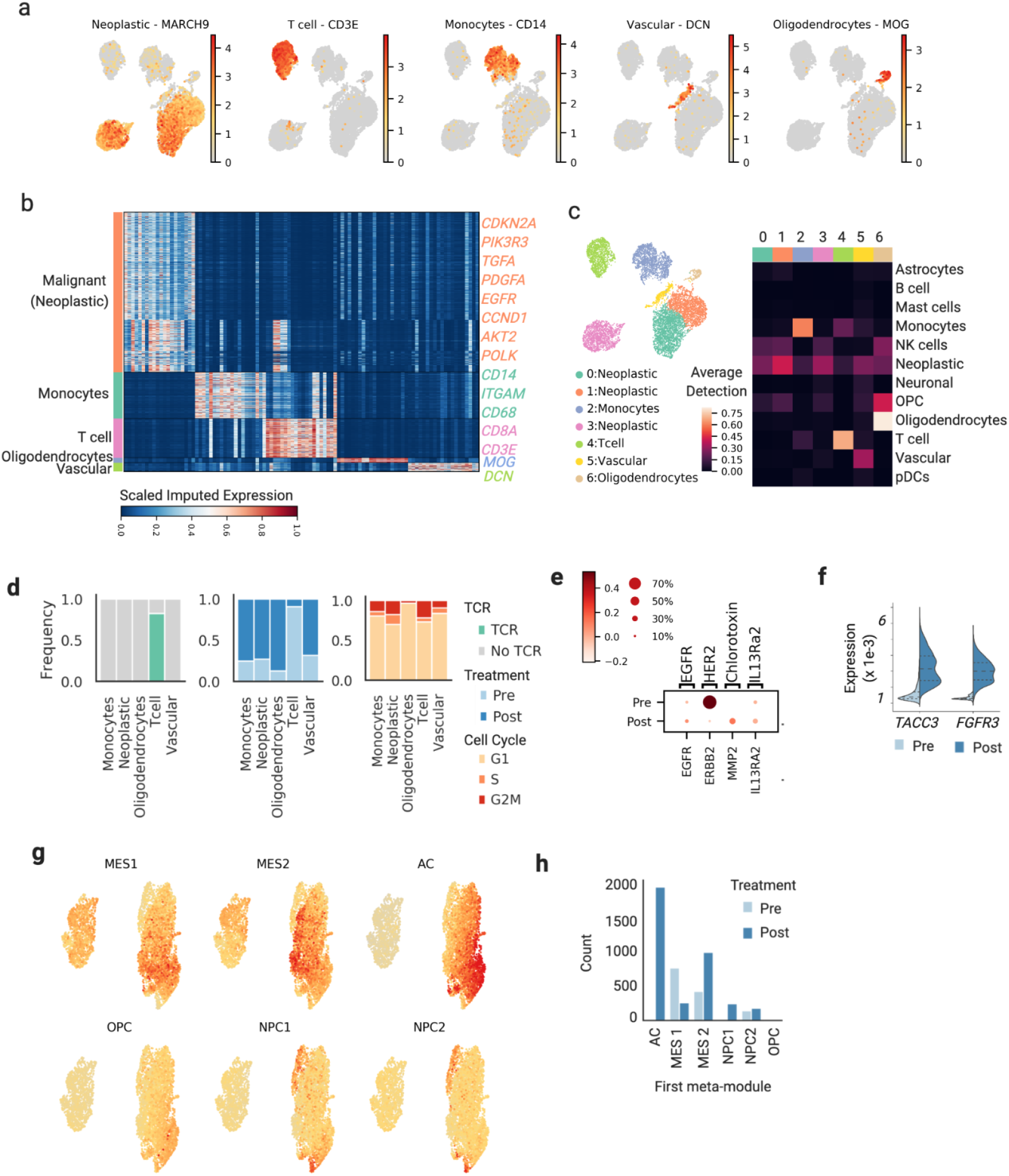
Tumor microenvironment characterization pre and post CAR T cell therapy. **a**, UMAP of all tumor resident cells colored by cell-type-specific markers. **b**, Heat map of differentially expressed genes (columns) between single cells belonging to inferred cell types, with canonical genes highlighted. **c**, UMAP of all GBM TME cells colored by original Leiden clustering and labeled by assigned cell type with highest detection score (left). Average gene detection per cluster in pre and post CAR T treated GBM tumors are based on previously published gene signatures of neuro and immune populations (right). **d**, Bar plots of relative cell proportions per cell type colored by TCR detection (left), treatment status (middle), and cell cycle phases (right). **e**, Dot plot of GBM surface antigen expression in malignant single cells pre- and post-treatment, with dots sized by percent expressing and colored by average gene expression. **f**, Violin plot of imputed single cell expression of oncogenic drivers. **g**, UMAP of all pre- and post-treatment malignant cells colored by gene module score corresponding to Mesenchymal/MES-like, Astrocytic/AS-like, Oligodendrocytic/OPC- like, and Neural progenitor/NPC-like GBM cellular states. **h**, Bar plot of cell count per treatment status with the highest score for each GBM meta-module.

**Extended Data Fig. 3.**
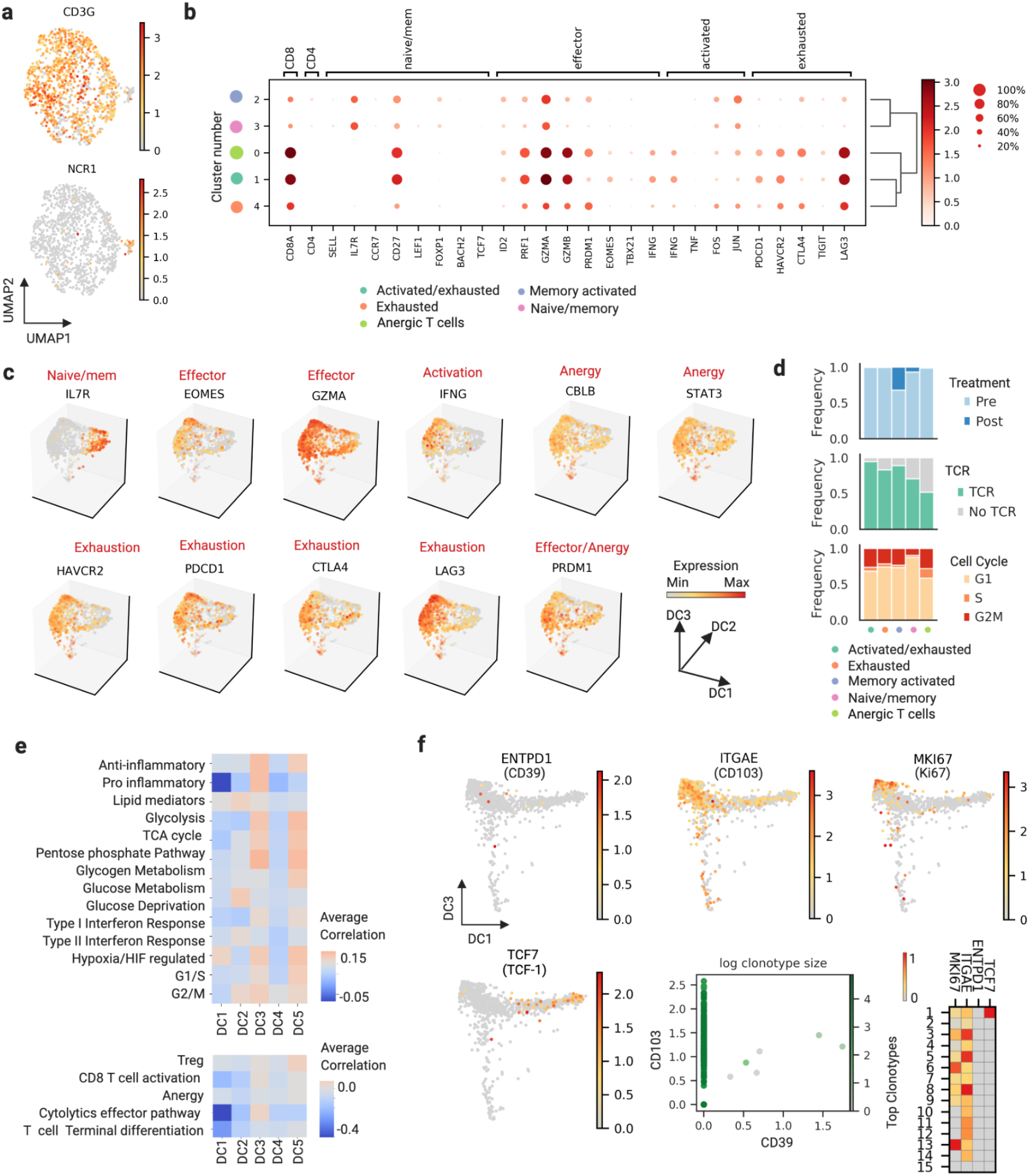
Characterization of T cell phenotypes present in pre and post CAR T cell treated tumors. **a**, UMAP of re-clustered T cells colored by T cell and NK cell marker gene expression identified small NK cell population. **b**, Dot plot of single cell gene expression of T cell phenotype marker genes in each T cell cluster. **c**, Diffusion map of tumor resident T cell populations using first three diffusion components colored by gene expression. **d**, Bar plots of relative cell proportions per T cell phenotypes colored by TCR detection (top), treatment status (middle), and cell cycle phases (bottom). **e**, Average correlation between diffusion component coordinate and gene expression of previously published inflammatory, metabolic, and T cell gene signatures for tumor resident T cells. **f**, Diffusion map (components 1 and 3) of tumor resident T cells colored by tumor-reactive genes (top) and self-renewal genes (bottom left). Scatterplot of CD39 and CD103 single cell co-expression painted by log clonotype size (bottom center). Matrix plot of average expression per top clonotype in pre- and post-treatment TILs, standardized per gene (bottom right).

**Extended Fig. 4.**
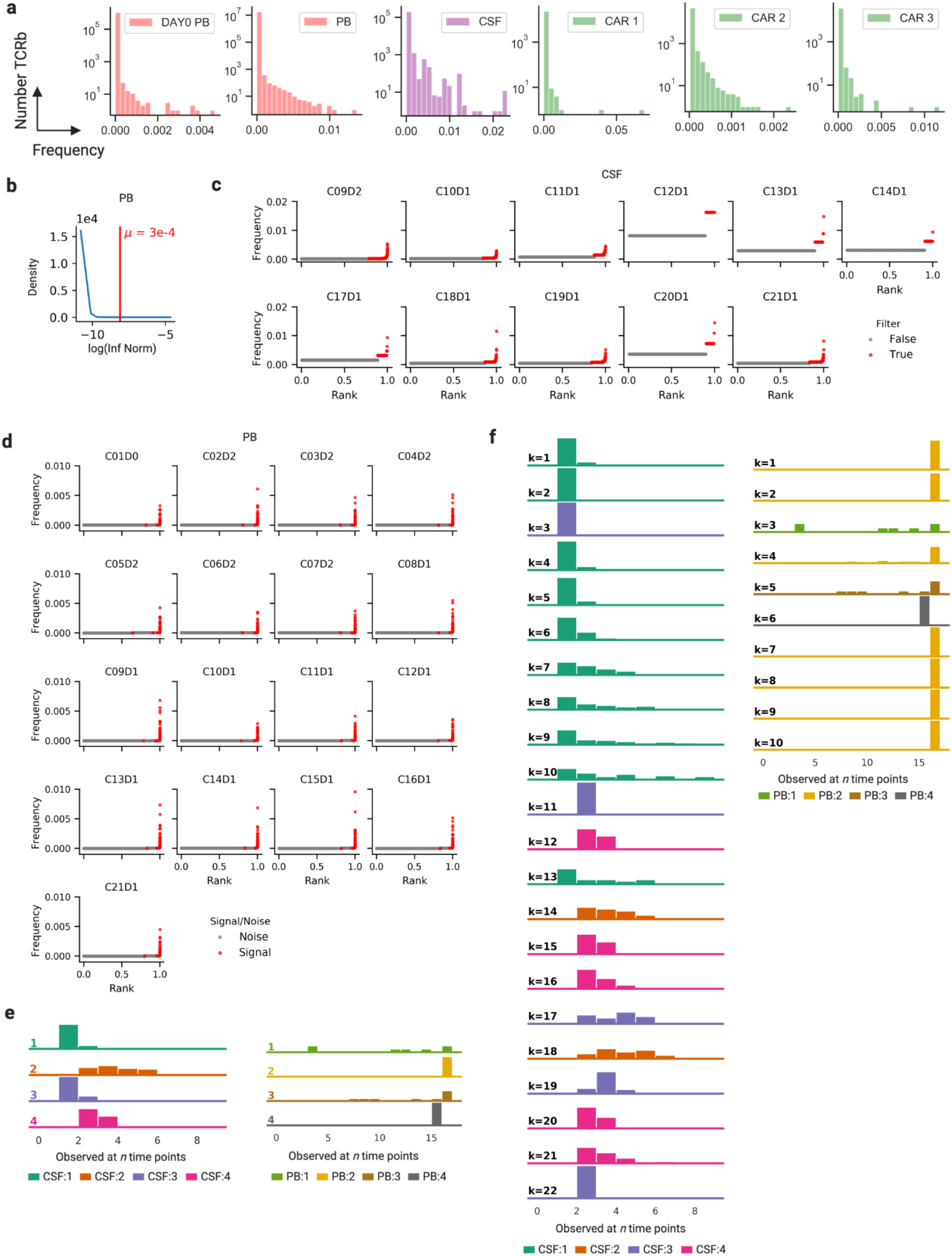
CSF and PB TRB dynamics signal - noise distribution. **a**, Histogram representing number of TRB in PB on Day 0, pre CAR T treatment, PB and CSF for each time point, and three CAR T products. **b**, Density plot of infinity norm (logarithmic scale) for PB TRB time series with signal-noise determination threshold μ marked (red). **c**, Scatter plot of CSF TRB time series rank vs. frequency at each time point colored by signal-noise determination filter. **d**, Scatter plot of PB TRB time series rank vs. frequency at each time point colored by signal-noise determination filter. **e,** Histograms calculated per k-shape cluster in CSF (left) and PB (right) of the number of time points at which the TRB is observed. **f,** Histograms calculated per k-means cluster in CSF (left) and PB (right) of the number of time points at which the TRB is observed. Histograms are colored by k-shape assignment.

**Extended Data Fig. 5.**
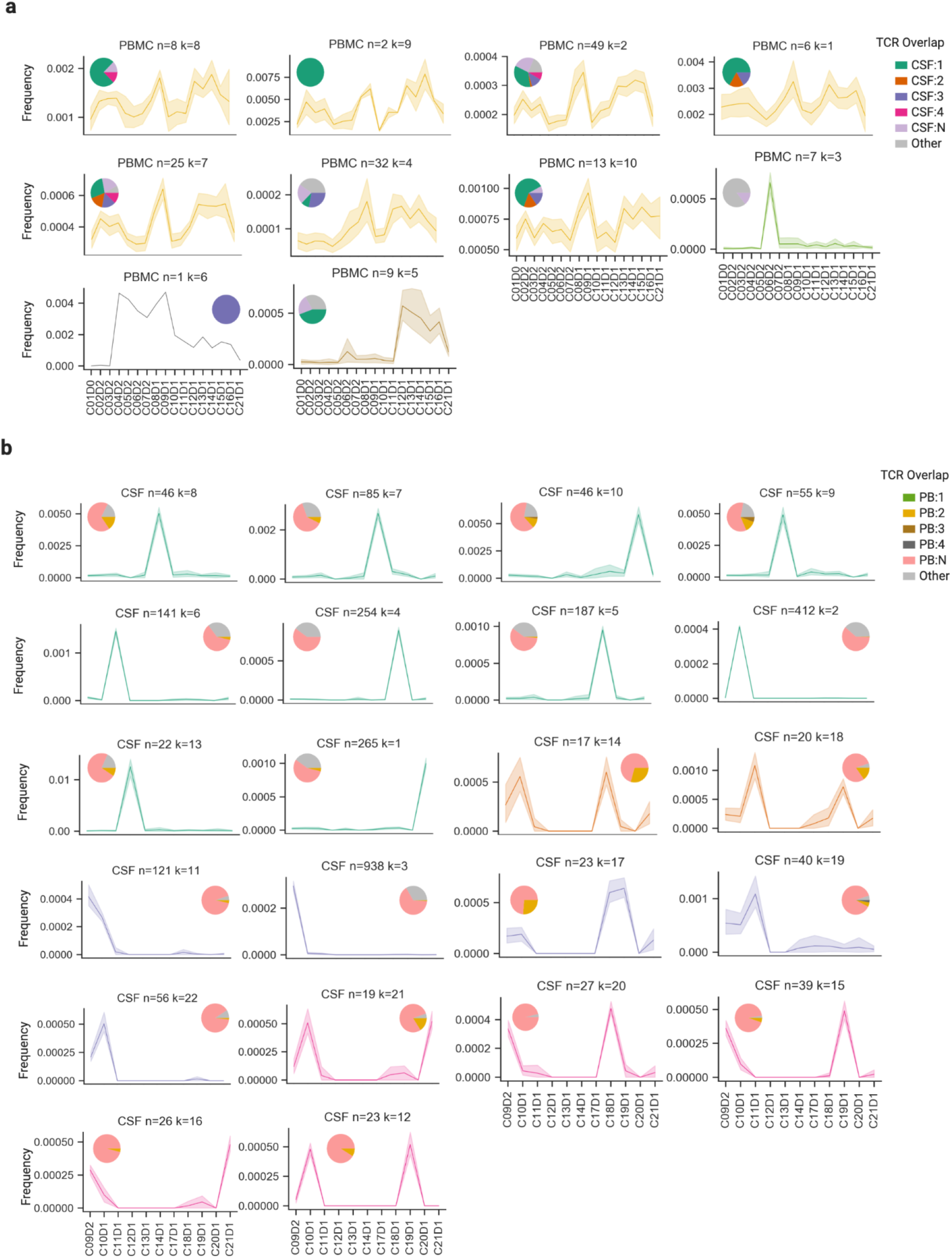
TRB clonal dynamics in peripheral blood (PB) and cerebrospinal fluid (CSF) measured by bulk TRB sequencing. **a**, Mean traces with bootstrapped 95% confidence interval of PB TRB time series grouped by k-means clustering and colored by k-shape signal dynamics. Pie charts indicate the fraction of TRB sequences from the cluster that overlaps with TRB sequences found in each of CSF noise or signal dynamics. **b,** Mean traces with bootstrapped 95% confidence interval of CSF TRB time series grouped by k-means cluster and colored by k-shape signal dynamics. Pie charts indicate the fraction of TRB sequences from the cluster that overlaps with TRB sequences found in each of PB noise or signal dynamics.

**Extended Data Fig. 6.**
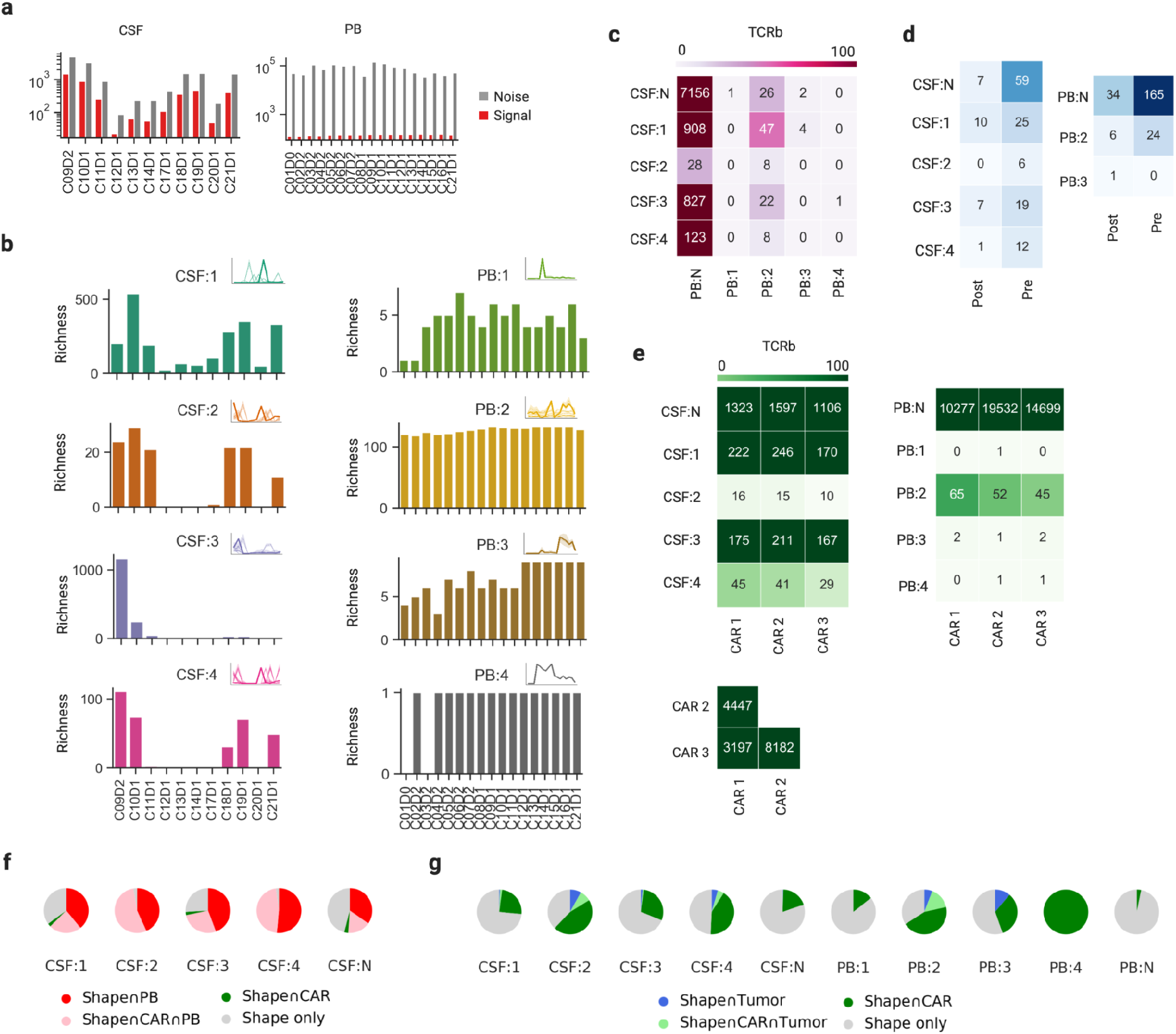
Tumor infiltrating lymphocytes (TILs) and CAR T cell product TRB richness overlap with CSF and PB TRB dynamics. **a**, Bar plot of CSF (left) and PB (right) of TRB richness per time point grouped by low frequency (Noise) or high frequency (Signal). **b**, Bar plot of TRB richness per time point for each signal dynamics in CSF (left) and PB (right). **c,** Heatmap of TRB richness of overlap between CSF and PB in noise and signal dynamics. **d**, Heatmap of TRB richness of overlap between TIL and CSF (left) or PB (right) noise and signal dynamics. **e,** Heatmap of TRB richness of overlap between each CAR T cell product and CSF (left) or PB (right) noise and signal dynamics, and overlap between the CAR T cell products (bottom). **f,** Pie charts of fraction of TRB sequences from each CSF noise and signal dynamics that overlaps with TRB sequences from PB, CAR T products, or both. **g,** Pie charts of fraction of TRB sequences from each CSF and PB noise and signal dynamics that overlaps with TRB sequences from tumor, CAR T products, or both.

**Extended Data Fig. 7.**
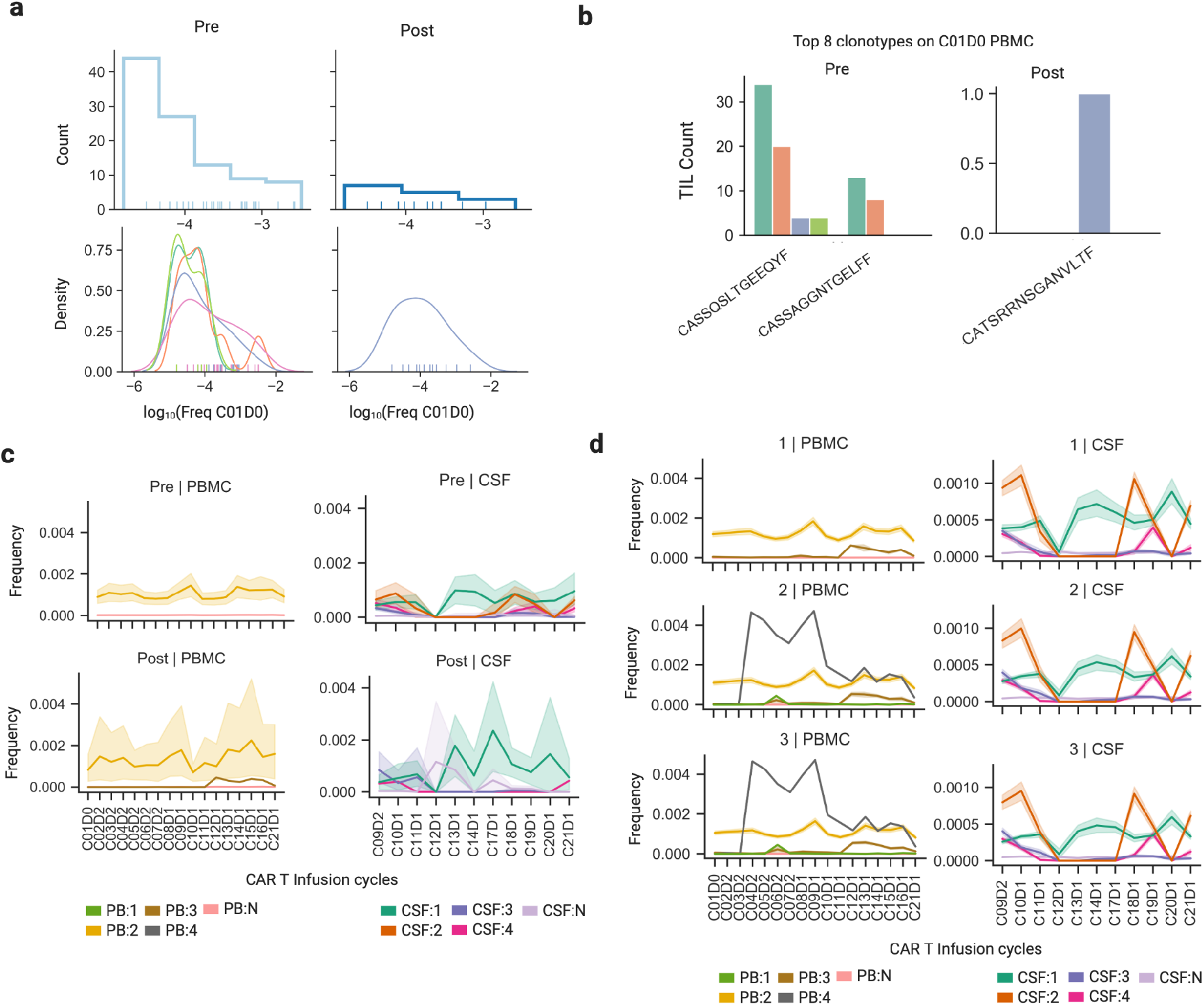
Tumor infiltrating lymphocytes (TILs) and CAR T cell TRB dynamics in with CSF and PB. **a**, Histogram of Pre (left) and Post (right) TIL overlap with PB C01D0, prior to CAR T cell therapy (top). Kernel density approximation of Pre and Post TIL count that overlap with PB C01D0, colored by TIL phenotype (bottom). **b,** Bar plot of cell counts of Pre (left) and Post (right) top 8 clonotypes that overlap with PB C01D0, colored by TIL phenotype. **c**, Mean traces with bootstrapped 95% confidence interval of PB (left) and CSF (right) TRB time series that overlap with TIL TRB sequences. **d**, Mean traces with bootstrapped 95% confidence interval of PB (left) and CSF (right) TRB time series that overlap with CAR T product 1, 2, and 3 (from top to bottom) TRB sequences.

## REFERENCES

Ayers, Mark, Jared Lunceford, Michael Nebozhyn, Erin Murphy, Andrey Loboda, David R. Kaufman, Andrew Albright, et al. 2017. “IFN-γ-Related mRNA Profile Predicts Clinical Response to PD-1 Blockade.” The Journal of Clinical Investigation 127 (8): 2930–40.

Azizi, Elham, Ambrose J. Carr, George Plitas, Andrew E. Cornish, Catherine Konopacki, Sandhya Prabhakaran, Juozas Nainys, et al. 2018. “Single-Cell Map of Diverse Immune Phenotypes in the Breast Tumor Microenvironment.” Cell 174 (5): 1293–1308.e36.

Beltra, Jean-Christophe, Sasikanth Manne, Mohamed S. Abdel-Hakeem, Makoto Kurachi, Josephine R. Giles, Zeyu Chen, Valentina Casella, et al. 2020. “Developmental Relationships of Four Exhausted CD8 T Cell Subsets Reveals Underlying Transcriptional and Epigenetic Landscape Control Mechanisms.” Immunity 52 (5): 825–41.e8.

Bray, Nicolas L., Harold Pimentel, Páll Melsted, and Lior Pachter. 2016. “Near-Optimal Probabilistic RNA-Seq Quantification.” Nature Biotechnology 34 (5): 525–27.

Brown, Christine E., Darya Alizadeh, Renate Starr, Lihong Weng, Jamie R. Wagner, Araceli Naranjo, Julie R. Ostberg, et al. 2016. “Regression of Glioblastoma after Chimeric Antigen Receptor T-Cell Therapy.” The New England Journal of Medicine 375 (26): 2561–69.

Brown, Christine E., Charles D. Warden, Renate Starr, Xutao Deng, Behnam Badie, Yate-Ching Yuan, Stephen J. Forman, and Michael E. Barish. 2013. “Glioma IL13Rα2 Is Associated with Mesenchymal Signature Gene Expression and Poor Patient Prognosis.” PloS One 8 (10): e77769.

Chen, Zeyu, Zhicheng Ji, Shin Foong Ngiow, Sasikanth Manne, Zhangying Cai, Alexander C. Huang, John Johnson, et al. 2019. “TCF-1-Centered Transcriptional Network Drives an Effector versus Exhausted CD8 T Cell-Fate Decision.” Immunity. https://doi.org/10.1016/j.immuni.2019.09.013.

Chu, Nathaniel D., Haixin Sarah Bi, Ryan O. Emerson, Anna M. Sherwood, Michael E. Birnbaum, Harlan S. Robins, and Eric J. Alm. 2019. “Longitudinal Immunosequencing in Healthy People Reveals Persistent T Cell Receptors Rich in Highly Public Receptors.” BMC Immunology. https://doi.org/10.1186/s12865-019-0300-5.

Darmanis, Spyros, Steven A. Sloan, Derek Croote, Marco Mignardi, Sophia Chernikova, Peyman Samghababi, Ye Zhang, et al. 2017. “Single-Cell RNA-Seq Analysis of Infiltrating Neoplastic Cells at the Migrating Front of Human Glioblastoma.” Cell Reports. https://doi.org/10.1016/j.celrep.2017.10.030.

Duhen, Thomas, Rebekka Duhen, Ryan Montler, Jake Moses, Tarsem Moudgil, Noel F. de Miranda, Cheri P. Goodall, et al. 2018. “Co-Expression of CD39 and CD103 Identifies Tumor-Reactive CD8 T Cells in Human Solid Tumors.” Nature Communications 9 (1): 2724.

Fang, Zhuoqing. n.d. GSEApy: Gene Set Enrichment Analysis in Python. Github. Accessed September 22, 2021. https://github.com/zqfang/GSEApy.

Gee, Marvin H., Arnold Han, Shane M. Lofgren, John F. Beausang, Juan L. Mendoza, Michael E. Birnbaum, Michael T. Bethune, et al. 2018. “Antigen Identification for Orphan T Cell Receptors Expressed on Tumor-Infiltrating Lymphocytes.” Cell 172 (3): 549–63.e16.

Haghverdi, Laleh, Florian Buettner, and Fabian J. Theis. 2015. “Diffusion Maps for High-Dimensional Single-Cell Analysis of Differentiation Data.” Bioinformatics 31 (18): 2989–98.

Halperin, Rebecca F., Winnie S. Liang, Sidharth Kulkarni, Erica E. Tassone, Jonathan Adkins, Daniel Enriquez, Nhan L. Tran, et al. 2019. “Leveraging Spatial Variation in Tumor Purity for Improved Somatic Variant Calling of Archival Tumor Only Samples.” Frontiers in Oncology. https://doi.org/10.3389/fonc.2019.00119.

Hegde, Meenakshi, Sujith K. Joseph, Farzana Pashankar, Christopher DeRenzo, Khaled Sanber, Shoba Navai, Tiara T. Byrd, et al. 2020. “Tumor Response and Endogenous Immune Reactivity after Administration of HER2 CAR T Cells in a Child with Metastatic Rhabdomyosarcoma.” Nature Communications. https://doi.org/10.1038/s41467-020-17175-8.

Hodges, Tiffany R., Martina Ott, Joanne Xiu, Zoran Gatalica, Jeff Swensen, Shouhao Zhou, Jason T. Huse, et al. 2017. “Mutational Burden, Immune Checkpoint Expression, and Mismatch Repair in Glioma: Implications for Immune Checkpoint Immunotherapy.” Neuro-Oncology 19 (8): 1047–57.

Huang, Huang, Chunlin Wang, Florian Rubelt, Thomas J. Scriba, and Mark M. Davis. 2020. “Analyzing the Mycobacterium Tuberculosis Immune Response by T-Cell Receptor Clustering with GLIPH2 and Genome-Wide Antigen Screening.” Nature Biotechnology 38 (10): 1194–1202.

Hunder, Naomi N., Herschel Wallen, Jianhong Cao, Deborah W. Hendricks, John Z. Reilly, Rebecca Rodmyre, Achim Jungbluth, Sacha Gnjatic, John A. Thompson, and Cassian Yee. 2008. “Treatment of Metastatic Melanoma with Autologous CD4+ T Cells against NY-ESO-1.” The New England Journal of Medicine 358 (25): 2698–2703.

Im, Se Jin, Masao Hashimoto, Michael Y. Gerner, Junghwa Lee, Haydn T. Kissick, Matheus C. Burger, Qiang Shan, et al. 2016. “Defining CD8 T Cells That Provide the Proliferative Burst after PD-1 Therapy.” Nature. https://doi.org/10.1038/nature19330.

Jeon, Myung-Shin, Alex Atfield, K. Venuprasad, Connie Krawczyk, Renu Sarao, Chris Elly, Chun Yang, et al. 2004. “Essential Role of the E3 Ubiquitin Ligase Cbl-B in T Cell Anergy Induction.” Immunity. https://doi.org/10.1016/j.immuni.2004.07.013.

Köster, Johannes, and Sven Rahmann. 2018. “Snakemake-a Scalable Bioinformatics Workflow Engine.” Bioinformatics 34 (20): 3600.

Lopez, Romain, Jeffrey Regier, Michael B. Cole, Michael I. Jordan, and Nir Yosef. 2018. “Deep Generative Modeling for Single-Cell Transcriptomics.” Nature Methods 15 (12): 1053–58.

Louveau, Antoine, Jasmin Herz, Maria Nordheim Alme, Andrea Francesca Salvador, Michael Q. Dong, Kenneth E. Viar, S. Grace Herod, et al. 2018. “CNS Lymphatic Drainage and Neuroinflammation Are Regulated by Meningeal Lymphatic Vasculature.” Nature Neuroscience 21 (10): 1380–91.

Louveau, Antoine, Igor Smirnov, Timothy J. Keyes, Jacob D. Eccles, Sherin J. Rouhani, J. David Peske, Noel C. Derecki, et al. 2015. “Structural and Functional Features of Central Nervous System Lymphatic Vessels.” Nature 523 (7560): 337–41.

Martin, Marcel. 2011. “Cutadapt Removes Adapter Sequences from High-Throughput Sequencing Reads.” EMBnet.journal. https://doi.org/10.14806/ej.17.1.200.

Melsted, Páll, Shannon Hateley, Isaac Charles Joseph, Harold Pimentel, Nicolas Bray, and Lior Pachter. n.d. “Fusion Detection and Quantification by Pseudoalignment.” https://doi.org/10.1101/166322.

Miller, Alexandra M., Ronak H. Shah, Elena I. Pentsova, Maryam Pourmaleki, Samuel Briggs, Natalie Distefano, Youyun Zheng, et al. 2019. “Tracking Tumour Evolution in Glioma through Liquid Biopsies of Cerebrospinal Fluid.” Nature 565 (7741): 654–58.

Neftel, Cyril, Julie Laffy, Mariella G. Filbin, Toshiro Hara, Marni E. Shore, Gilbert J. Rahme, Alyssa R. Richman, et al. 2019. “An Integrative Model of Cellular States, Plasticity, and Genetics for Glioblastoma.” Cell 178 (4): 835–49.e21.

Pimentel, Harold, Nicolas L. Bray, Suzette Puente, Páll Melsted, and Lior Pachter. 2017. “Differential Analysis of RNA-Seq Incorporating Quantification Uncertainty.” Nature Methods 14 (7): 687–90.

Sahoo, Prativa, Ram K. S. Rathore, Rishi Awasthi, Bhaswati Roy, Sanjay Verma, Divya Rathore, Sanjay Behari, et al. 2013. “Subcompartmentalization of Extracellular Extravascular Space (EES) into Permeability and Leaky Space with Local Arterial Input Function (AIF) Results in Improved Discrimination between High- and Low-Grade Glioma Using Dynamic Contrast-Enhanced (DCE) MRI.” Journal of Magnetic Resonance Imaging: JMRI 38 (3): 677–88.

Salou, Marion, Alexandra Garcia, Laure Michel, Anne Gainche-Salmon, Delphine Loussouarn, Bryan Nicol, Flora Guillot, et al. 2015. “Expanded CD 8 T-cell Sharing between Periphery and CNS in Multiple Sclerosis.” Annals of Clinical and Translational Neurology. https://doi.org/10.1002/acn3.199.

Schäfer, Niklas, Gerrit H. Gielen, Laurèl Rauschenbach, Sied Kebir, Andreas Till, Roman Reinartz, Matthias Simon, et al. 2019. “Longitudinal Heterogeneity in Glioblastoma: Moving Targets in Recurrent versus Primary Tumors.” Journal of Translational Medicine 17 (1): 96.

Scheper, Wouter, Sander Kelderman, Lorenzo F. Fanchi, Carsten Linnemann, Gavin Bendle, Marije A. J. de Rooij, Christian Hirt, et al. 2019. “Low and Variable Tumor Reactivity of the Intratumoral TCR Repertoire in Human Cancers.” Nature Medicine 25 (1): 89–94.

Siddiqui, Imran, Karin Schaeuble, Vijaykumar Chennupati, Silvia A. Fuertes Marraco, Sandra Calderon-Copete, Daniela Pais Ferreira, Santiago J. Carmona, et al. 2019. “Intratumoral Tcf1PD-1CD8 T Cells with Stem-like Properties Promote Tumor Control in Response to Vaccination and Checkpoint Blockade Immunotherapy.” Immunity 50 (1): 195–211.e10.

Singh, Devendra, Joseph Minhow Chan, Pietro Zoppoli, Francesco Niola, Ryan Sullivan, Angelica Castano, Eric Minwei Liu, et al. 2012. “Transforming Fusions of FGFR and TACC Genes in Human Glioblastoma.” Science 337 (6099): 1231–35.

Song, Eric, Tianyang Mao, Huiping Dong, Ligia Simoes Braga Boisserand, Salli Antila, Marcus Bosenberg, Kari Alitalo, Jean-Leon Thomas, and Akiko Iwasaki. 2020. “VEGF-C-Driven Lymphatic Drainage Enables Immunosurveillance of Brain Tumours.” Nature 577 (7792): 689–94.

Tirosh, Itay, Benjamin Izar, Sanjay M. Prakadan, Marc H. Wadsworth 2nd, Daniel Treacy, John J. Trombetta, Asaf Rotem, et al. 2016. “Dissecting the Multicellular Ecosystem of Metastatic Melanoma by Single-Cell RNA-Seq.” Science 352 (6282): 189–96.

Verhaak, Roel G. W., Katherine A. Hoadley, Elizabeth Purdom, Victoria Wang, Yuan Qi, Matthew D. Wilkerson, C. Ryan Miller, et al. 2010. “Integrated Genomic Analysis Identifies Clinically Relevant Subtypes of Glioblastoma Characterized by Abnormalities in PDGFRA, IDH1, EGFR, and NF1.” Cancer Cell 17 (1): 98–110.

Virtanen, Pauli, Ralf Gommers, Travis E. Oliphant, Matt Haberland, Tyler Reddy, David Cournapeau, Evgeni Burovski, et al. 2020. “SciPy 1.0: Fundamental Algorithms for Scientific Computing in Python.” Nature Methods 17 (3): 261–72.

Wolf, F. Alexander, Philipp Angerer, and Fabian J. Theis. 2018. “SCANPY: Large-Scale Single-Cell Gene Expression Data Analysis.” Genome Biology 19 (1): 15.

Yi, Lynn, Harold Pimentel, Nicolas L. Bray, and Lior Pachter. 2018. “Gene-Level Differential Analysis at Transcript-Level Resolution.” Genome Biology 19 (1): 53.

Yost, Kathryn E., Ansuman T. Satpathy, Daniel K. Wells, Yanyan Qi, Chunlin Wang, Robin Kageyama, Katherine L. McNamara, et al. 2019. “Clonal Replacement of Tumor-Specific T Cells Following PD-1 Blockade.” Nature Medicine 25 (8): 1251–59.

